# Limitations of principal components in quantitative genetic association models for human studies

**DOI:** 10.1101/2022.03.25.485885

**Authors:** Yiqi Yao, Alejandro Ochoa

**Affiliations:** Department of Biostatistics and Bioinformatics, Duke University, Durham, NC 27705, USA; Duke Center for Statistical Genetics and Genomics, Duke University, Durham, NC 27705, USA; BenHealth Consulting, Shanghai, Shanghai, 200023, China

## Abstract

Principal Component Analysis (PCA) and the Linear Mixed-effects Model (LMM), sometimes in combination, are the most common genetic association models. Previous PCA-LMM comparisons give mixed results, unclear guidance, and have several limitations, including not varying the number of principal components (PCs), simulating simple population structures, and inconsistent use of real data and power evaluations. We evaluate PCA and LMM both varying number of PCs in realistic genotype and complex trait simulations including admixed families, subpopulation trees, and real multiethnic human datasets with simulated traits. We find that LMM without PCs usually performs best, with the largest effects in family simulations and real human datasets and traits without environment effects. Poor PCA performance on human datasets is driven by large numbers of distant relatives more than the smaller number of closer relatives. While PCA was known to fail on family data, we report strong effects of family relatedness in genetically diverse human datasets, not avoided by pruning close relatives. Environment effects driven by geography and ethnicity are better modeled with LMM including those labels instead of PCs. This work better characterizes the severe limitations of PCA compared to LMM in modeling the complex relatedness structures of multiethnic human data for association studies.

## 1 Introduction

The goal of a genetic association study is to identify loci whose genotype variation is significantly correlated to given trait. Naive association tests assume that genotypes are drawn independently from a common allele frequency. This assumption does not hold for structured populations, which includes multiethnic cohorts and admixed individuals (ancient relatedness), and for family data (recent relatedness) [1]. Association studies of admixed and multiethnic cohorts, the focus of this work, are becoming more common, are believed to be more powerful, and are necessary to bring more equity to genetic medicine [2–22]. When insufficient approaches are applied to data with relatedness, their association statistics are miscalibrated, resulting in excess false positives and loss of power [1, 23, 24]. Therefore, many specialized approaches have been developed for genetic association under relatedness, of which PCA and LMM are the most popular.

Genetic association with PCA consists of including the top eigenvectors of the population kinship matrix as covariates in a generalized linear model [25–27]. These top eigenvectors are a new set of coordinates for individuals that are commonly referred to as PCs in genetics [28], the convention adopted here, but in other fields PCs instead denote what in genetics would be the projections of loci onto eigenvectors, which are new independent coordinates for loci [29]. The direct ancestor of PCA association is structured association, in which inferred ancestry (genetic cluster membership, often corresponding with labels such as “European”, “African”, “Asian”, etc.) or admixture proportions of these ancestries are used as regression covariates [30]. These models are deeply connected because PCs map to ancestry empirically [31, 32] and theoretically [33–36], and they work as well as global ancestry in association studies but are estimated more easily [27, 28, 31, 37]. Another approach closely related to PCA is nonmetric multidimensional scaling [38]. PCs are also proposed for modeling environment effects that are correlated to ancestry, for example, through geography [20, 39, 40]. The strength of PCA is its simplicity, which as covariates can be readily included in more complex models, such as haplotype association [41] and polygenic models [42]. However,

PCA assumes that the underlying relatedness space is low dimensional (or low rank), so it can be well modeled with a small number of PCs, which may limit its applicability. PCA is known to be inadequate for family data [28, 38, 43, 44], which is called “cryptic relatedness” when it is unknown to the researchers, but no other troublesome cases have been confidently identified. Recent work has focused on developing more scalable versions of the PCA algorithm [45–49]. PCA remains a popular and powerful approach for association studies.

The other dominant association model under relatedness is the LMM, which includes a random effect parameterized by the kinship matrix. Unlike PCA, LMM does not assume that relatedness is low-dimensional, and explicitly models families via the kinship matrix. Early LMMs used kinship matrices estimated from known pedigrees or using methods that captured recent relatedness only, and modeled population structure (ancestry) as fixed effects [37, 38, 50]. Modern LMMs estimate kinship from genotypes using a non-parametric estimator, often referred to as a genetic relationship matrix, that captures the combined covariance due to family relatedness and ancestry [1, 51, 52]. Like PCA, LMM has also been proposed for modeling environment correlated to genetics [53, 54]. The classic LMM assumes a quantitative (continuous) complex trait, the focus of our work. Although case-control (binary) traits and their underlying ascertainment are theoretically a challenge [55], LMMs have been applied successfully to balanced case-control studies [1, 56] and simulations [44, 57, 58], and have been adapted for unbalanced case-control studies [59]. However, LMMs tend to be considerably slower than PCA and other models, so much effort has focused on improving their runtime and scalability [51, 56, 59–67].

An LMM variant that incorporates PCs as fixed covariates is tested thoroughly in our work. Since PCs are the top eigenvectors of the same kinship matrix estimate used in modern LMMs [1, 3, 40, 68], then population structure is modeled twice in an LMM with PCs. However, some previous work has found the apparent redundancy of an LMM with PCs beneficial [40, 44, 69], while others did not [68, 70], and the approach continues to be used [71, 72] though not always [73]. (Recall that early LMMs used kinship to model family relatedness only, so population structure had to be modeled separately in those models, in practice as admixture fractions instead of PCs [37, 38, 50].) The LMM with PCs (vs no PCs) is also believed to help better model loci that have experienced selection [44, 53] and environment effects correlated with genetics [40].

LMM and PCA are closely related models [1, 3, 40, 68], so similar performance is expected particularly under low-dimensional relatedness. Direct comparisons have yielded mixed results, with several studies finding superior performance for LMM, notably from papers promoting advances in LMMs, while many others report comparable performance (Table 1). No papers find that PCA outperforms LMM decisively, although PCA occasionally performs better in isolated and artificial cases or individual measures, often with unknown significance. Previous studies generally used either only simulated or only real genotypes, with only two studies using both. The simulated genotype studies, which tended to have low model dimensions and *F*_ST_, were more likely to report ties or mixed results (6/8), whereas real genotypes tended to clearly favor LMMs (9/11). Similarly, 10/12 papers with quantitative traits favor LMMs, whereas 6/9 papers with case-control traits gave ties or mixed results—the only factor we do not explore in this work. Additionally, although all previous evaluations measured type I error (or proxies such as genomic inflation factors [23] or QQ plots), a large fraction (6/17) did not measure power (or proxies such as ROC curves), and only four used more than one number of PCs for PCA. Lastly, no consensus has emerged as to why LMM might outperform PCA or vice versa [3, 44, 58, 78], or which features of the real datasets are critical for the LMM advantage other than family relatedness, resulting in unclear guidance for using PCA. Hence, our work includes real and simulated genotypes with higher model dimensions and *F*_ST_ matching that of multiethnic human cohorts [52, 79], we vary the number of PCs, and measure robust proxies for type I error control and calibrated power.

**Table 1:**
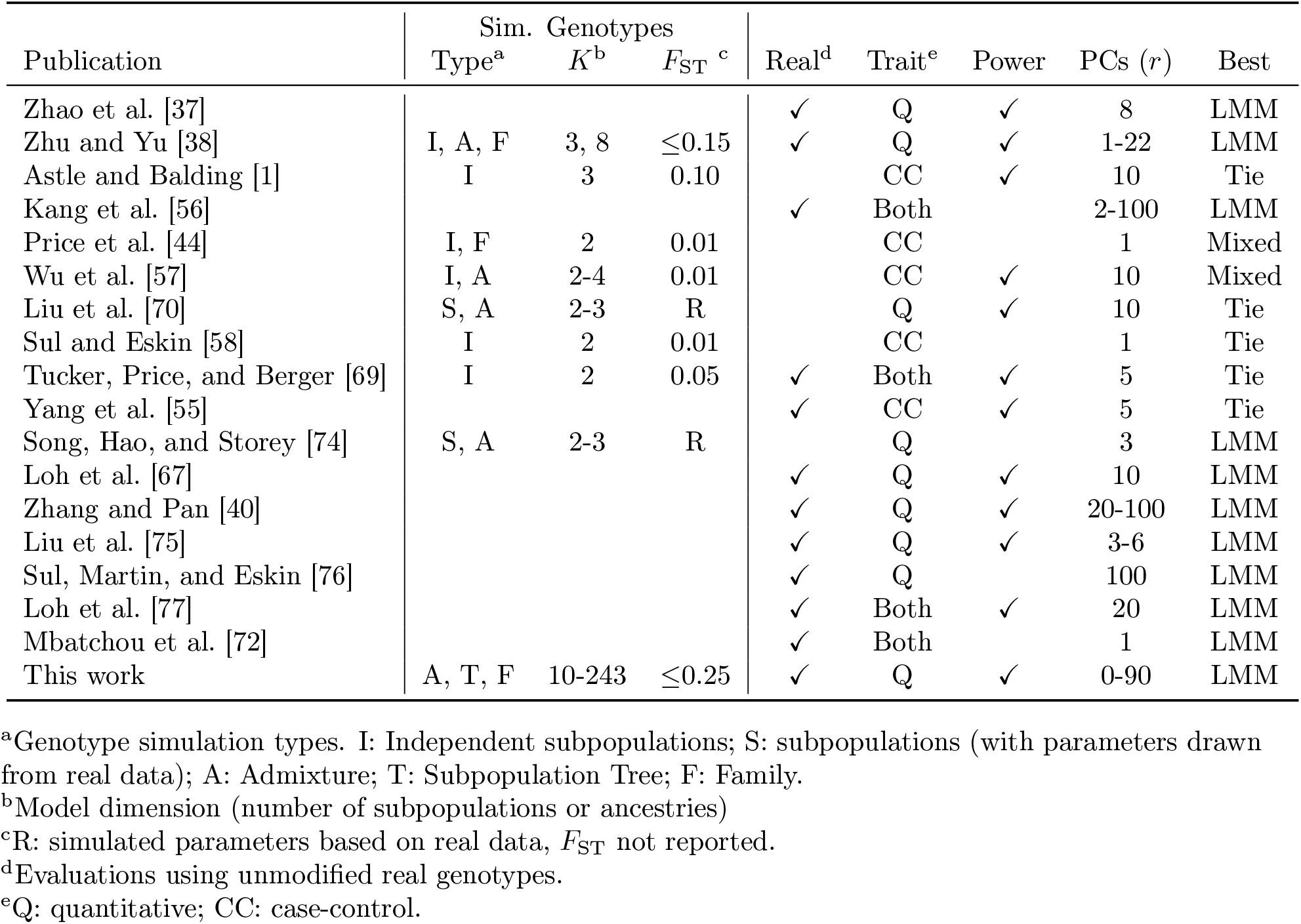
Previous PCA-LMM evaluations in the literature.

In this work, we evaluate the PCA and LMM association models under various numbers of PCs, which are included in LMMs too. We use genotype simulations (admixture, family, and subpopulation tree models) and three real datasets: the 1000 Genomes Project [80, 81], the Human Genome Diversity Panel (HGDP) [82–84], and Human Origins [85–88]. We simulate quantitative traits from two models: fixed effect sizes (FES) construct coefficients inverse to allele frequency, which matches real data [71, 89, 90] and corresponds to high pleiotropy and strong balancing selection [91] and strong negative selection [71, 90], which are appropriate assumptions for diseases; and random coefficients (RC), which are drawn independent of allele frequency, and corresponds to neutral traits [71, 91]. LMM without PCs consistently performs best in simulations without environment, and greatly outperforms PCA in the family simulation and in all real datasets. The tree simulations, which model subpopulations with the tree but exclude family structure, do not recapitulate the real data results, suggesting that family relatedness in real data is the reason for poor PCA performance. Lastly, removing up to 4th degree relatives in the real datasets recapitulates poor PCA performance, showing that the more numerous distant relatives explain the result, and suggesting that PCA is generally not an appropriate model for real data. We find that both LMM and PCA are able to model environment effects correlated with genetics, and LMM with PCs gains a small advantage in this setting only, but direct modeling of environment performs much better. All together, we find that LMMs without PCs are generally a preferable association model, and present novel simulation and evaluation approaches to measure the performance of these and other genetic association approaches.

## 2 Materials and Methods

### 2.1 The complex trait model and PCA and LMM approximations

Let *x*_*ij*_ ∈ {0, 1, 2} be the genotype at the biallelic locus *i* for individual *j*, which counts the number of reference alleles. Suppose there are *n* individuals and *m* loci, **X** = (*x*_*ij*_) is their *m × n* genotype matrix, and **y** is the length-*n* column vector of individual trait values. The additive linear model for a quantitative (continuous) trait is:

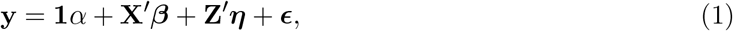

where **1** is a length-*n* vector of ones, *α* is the scalar intercept coefficient, ***β*** is the length-*m* vector of locus coefficients, **Z** is a design matrix of environment effects and other covariates, ***η*** is the vector of environment coefficients, ***ϵ*** is a length-*n* vector of residuals, and the prime symbol (*′*) denotes matrix transposition. The residuals follow *ϵ*_*j*_ ∼ Normal(0, 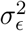) independently per individual *j*, for some 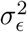.

The full model of Eq. (1), which has a coefficient for each of the *m* loci, is underdetermined in current datasets where *m* ≫ *n*. The PCA and LMM models, respectively, approximate the full model fit at a single locus *i*:

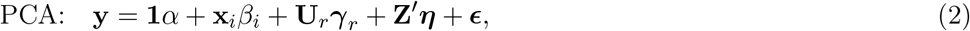

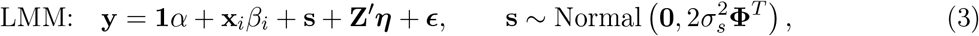

where **x**_*i*_ is the length-*n* vector of genotypes at locus *i* only, *β*_*i*_ is the locus coefficient, **U**_*r*_ is an *n* × *r* matrix of PCs, ***γ***_*r*_ is the length-*r* vector of PC coefficients, **s** is a length-*n* vector of random effects, 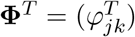 is the *n* × *n* kinship matrix conditioned on the ancestral population *T*, and 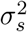 is a variance factor. Both models condition the regression of the focal locus *i* on an approximation of the total polygenic effect **X**′***β*** with the same covariance structure, which is parameterized by the kinship matrix. Under the kinship model, genotypes are random variables obeying

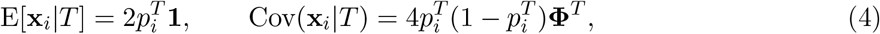

where 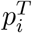 is the ancestral allele frequency of locus *i* [1, 92–94]. Assuming independent loci, the covariance of the polygenic effect is

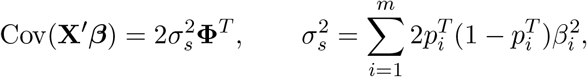

which is readily modeled by the LMM random effect **s**, where the difference in mean is absorbed by the intercept. Alternatively, consider the eigendecomposition of the kinship matrix **Φ**^*T*^ = **UΛU**′ where **U** is the *n* × *n* eigenvector matrix and **Λ** is the *n* × *n* diagonal matrix of eigenvalues. The random effect can be written as

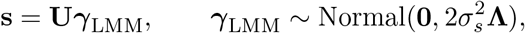

which follows from the affine transformation property of multivariate normal distributions. Therefore, the PCA term **U**_*r*_***γ***_*r*_ can be derived from the above equation under the additional assumption that the kinship matrix has approximate rank *r* and the coefficients ***γ***_*r*_ are fit without constraints. In contrast, the LMM uses all eigenvectors, while effectively shrinking their coefficients ***γ***_LMM_ as all random effects models do, although these parameters are marginalized [1, 3, 40, 68]. PCA has more parameters than LMM, so it may overfit more: ignoring the shared terms in Eqs. (2) and (3), PCA fits *r* parameters (length of ***γ***), whereas LMMs fit only one 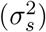.

In practice, the kinship matrix used for PCA and LMM is estimated with variations of a methodof-moments formula applied to standardized genotypes **X**_*S*_, which is derived from Eq. (4):

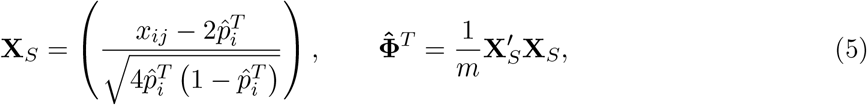

where the unknown 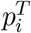 is estimated by 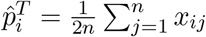 [26, 51, 55, 56, 59, 63, 65, 67, 76]. However, this kinship estimator has a complex bias that differs for every individual pair, which arises due to the use of this estimated 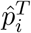 [52, 79]. Nevertheless, in PCA and LMM these biased estimates perform as well as unbiased ones [95].

We selected fast and robust software implementing the basic PCA and LMM models. PCA association was performed with `plink2` [96]. The quantitative trait association model is a linear regression with covariates, evaluated using the t-test. PCs were calculated with `plink2`, which equal the top eigenvectors of Eq. (5) after removing loci with minor allele frequency MAF < 0.1.

LMM association was performed using GCTA [55, 63]. Its kinship estimator equals Eq. (5). PCs were calculated using GCTA from its kinship estimate. Association significance is evaluated with a score test. In the small simulation only, GCTA with large numbers of PCs had convergence and singularity errors in some replicates, which were treated as missing data.

### 2.2 Simulations

Every simulation was replicated 50 times, drawing anew all genotypes (except for real datasets) and traits. Below we use the notation 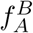 for the inbreeding coefficient of a subpopulation *A* from another subpopulation *B* ancestral to *A*. In the special case of the *total* inbreeding of *A*, 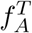, *T* is an overall ancestral population, which is ancestral to every individual under consideration, such as the most recent common ancestor (MRCA) population.

#### 2.2.1 Genotype simulation from the admixture model

The basic admixture model is as described previously [52] and is implemented in the R package `bnpsd`. Both Large and Family simulations have *n* = 1, 000 individuals, while Small has *n* = 100. The number of loci is *m* = 100, 000. Individuals are admixed from *K* = 10 intermediate subpopulations, or ancestries. Each subpopulation *S*_*u*_ (*u* ∈ {1, …, *K*}) is at coordinate *u* and has an inbreeding coefficient 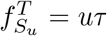 for some *τ*. Ancestry proportions *q*_*ju*_ for individual *j* and *S*_*u*_ arise from a random walk with spread *σ* on the 1D geography, and *τ* and *σ* are fit to give *F*_ST_ = 0.1 and mean kinship 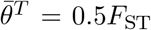 for the admixed individuals [52]. Random ancestral allele frequencies 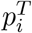, subpopulation allele frequencies 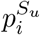, individual-specific allele frequencies *π*_*ij*_, and genotypes *x*_*ij*_ are drawn from this hierarchical model:

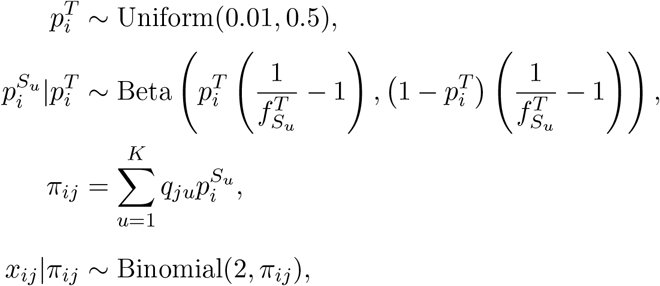

where this Beta is the Balding-Nichols distribution [97] with mean 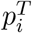 and variance 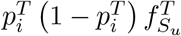. Fixed loci (*i* where *x*_*ij*_ = 0 for all *j*, or *x*_*ij*_ = 2 for all *j*) are drawn again from the model, starting from 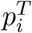, iterating until no loci are fixed. Each replicate draws a genotypes starting from 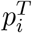.

As a brief aside, we prove that global ancestry proportions as covariates is equivalent in expectation to using PCs under the admixture model. Note that the latent space of **X**, which is the subspace to which the data is constrained by the admixture model, is given by (*π*_*ij*_), which has *K* dimensions (number of columns of **Q** = (*q*_*ju*_)), so the top *K* PCs span this space. Since associations include an intercept term (**1***α* in Eq. (2)), estimated PCs are orthogonal to **1** (note 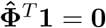 because **X**_*S*_**1** = **0**), and the sum of rows of **Q** sums to one, then only *K* − 1 PCs plus the intercept are needed to span the latent space of this admixture model.

#### 2.2.2 Genotype simulation from random admixed families

We simulated a pedigree with admixed founders, no close relative pairings, assortative mating based on a 1D geography (to preserve admixture structure), random family sizes, and arbitrary numbers of generations (20 here). This simulation is implemented in the R package `simfam`. Generations are drawn iteratively. Generation 1 has *n* = 1000 individuals from the above admixture simulation ordered by their 1D geography. Local kinship measures pedigree relatedness; in the first generation, everybody is locally unrelated and outbred. Individuals are randomly assigned sex. In the next generation, individuals are paired iteratively, removing random males from the pool of available males and pairing them with the nearest available female with local kinship < 1*/*4^3^ (stay unpaired if there are no matches), until there are no more available males or females. Let *n* = 1000 be the desired population size, *n*_*m*_ = 1 the minimum number of children per family and *n*_*f*_ the number of families (paired parents) in the current generation, then the number of additional children (beyond the minimum) is drawn from Poisson(*n/n*_*f*_ − *n*_*m*_). Let *δ* be the difference between desired and current population sizes. If *δ* > 0, then *δ* random families are incremented by 1. If *δ* < 0, then |*δ*| random families with at least *n*_*m*_ + 1 children are decremented by 1. If |*δ*| exceeds the number of families, all families are incremented or decremented as needed and the process is iterated. Children are assigned sex randomly, and are reordered by the average coordinate of their parents. Children draw alleles from their parents independently per locus. A new random pedigree is drawn for each replicate, as well as new founder genotypes from the admixture model.

#### 2.2.3 Genotype simulation from a subpopulation tree model

This model draws subpopulations allele frequencies from a hierarchical model parameterized by a tree, which is also implemented in `bnpsd` and relies on the R package `ape` for general tree data structures and methods [98]. The ancestral population *T* is the root, and each node is a subpopulation *S*_*w*_ indexed arbitrarily. Each edge between *S*_*w*_ and its parent population *P*_*w*_ has an inbreeding coefficient 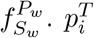 are drawn from a given distribution, which is constructed to mimic each real dataset in Appendix A. Given the allele frequencies 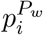 of the parent population, *S*_*w*_’s allele frequencies are drawn from:

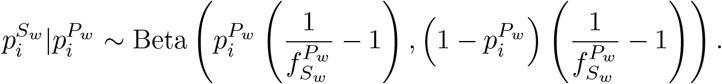

Individuals *j* in *S*_*w*_ draw genotypes from its allele frequency: 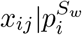 ∼ Binomial 2, 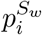). Loci with MAF < 0.01 are drawn again starting from the 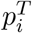 distribution, iterating until no such loci remain.

#### 2.2.4 Fitting subpopulation tree to real data

We developed new methods to fit trees to real data based on unbiased kinship estimates from `popkin`, implemented in `bnpsd`. A tree with given inbreeding coefficients 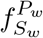 for its edges (between subpopulation *S*_*w*_ and its parent *P*_*w*_) gives rise to a coancestry matrix 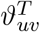 for a subpopulation pair (*S*_*u*_, *S*_*v*_), and the goal is to recover these edge inbreeding coefficients from coancestry estimates. Coancestry values are total inbreeding coefficients of the MRCA population of each subpopulation pair. Therefore, we calculate 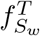 for every *S*_*w*_ recursively from the root as follows. Nodes with parent *P*_*w*_ = *T* are already as desired. Given 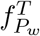, the desired 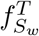 is calculated via the “additive edge” *δ*_*w*_ [52]:

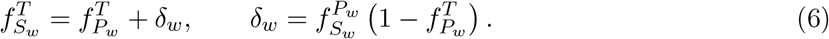

These *δ*_*w*_ ≥ 0 because 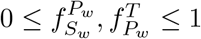 for every *w*. Edge inbreeding coefficients can be recovered from additive edges: 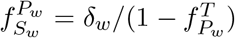. Overall, coancestry values are sums of *δ*_*w*_ over common ancestor nodes,

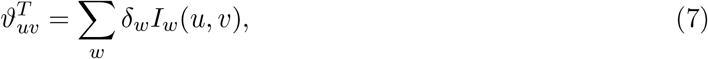

where the sum includes all *w*, and *I*_*w*_(*u, v*) equals 1 if *S*_*w*_ is a common ancestor of *S*_*u*_, *S*_*v*_, 0 otherwise. Note that *I*_*w*_(*u, v*) reflects tree topology and *δ*_*w*_ edge values.

To estimate population-level coancestry, first kinship 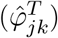 is estimated using `popkin` [52]. Individual coancestry 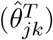 is estimated from kinship using

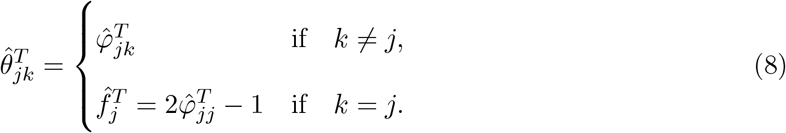

Lastly, coancestry 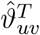 between subpopulations are averages of individual coancestry values:

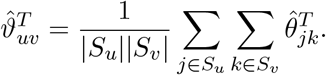

Topology is estimated with hierarchical clustering using the weighted pair group method with arithmetic mean [99], with distance function 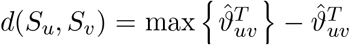, which succeeds due to the monotonic relationship between node depth and coancestry (Eq. (7)). This algorithm recovers the true topology from the true coancestry values, and performs well for estimates from genotypes.

To estimate tree edge lengths, first *δ*_*w*_ are estimated from 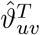 and the topology using Eq. (7) and non-negative least squares linear regression [100] (implemented in `nnls` [101]) to yield non-negative *δ*_*w*_, and 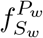 are calculated from *δ*_*w*_ by reversing Eq. (6). To account for small biases in coancestry estimation, an intercept term *δ*_0_ is included (*I*_0_(*u, v*) = 1 for all *u, v*), and when converting *δ*_*w*_ to 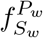, *δ*_0_ is treated as an additional edge to the root, but is ignored when drawing allele frequencies from the tree.

#### 2.2.5 Trait Simulation

Traits are simulated from the quantitative trait model of Eq. (1), with novel bias corrections for simulating the desired heritability from real data relying on the unbiased kinship estimator `popkin` [52]. This simulation is implemented in the R package `simtrait`. All simulations have a fixed narrow-sense heritability of *h*^2^, a variance proportion due to environment effects 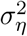, and residuals are drawn from *ϵ*_*j*_ ∼ Normal(0, 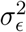) with 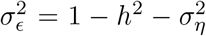. The number of causal loci *m*_1_, which determines the average coefficient size, is chosen with the heuristic formula *m*_1_ = round(*nh*^2^*/*8), which empirically balances power well with varying *n* and *h*^2^. The set of causal loci *C* is drawn anew for each replicate, from loci with MAF ≥ 0.01 to avoid rare causal variants, which are not discoverable by PCA or LMM at the sample sizes we considered. Letting 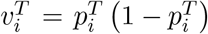, the effect size of locus *i* equals 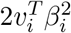, its contribution of the trait variance [102]. Under the *fixed effect sizes* (FES) model, initial causal coefficients are

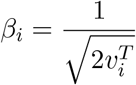

for known 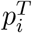; otherwise 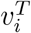 is replaced by the unbiased estimator [52] 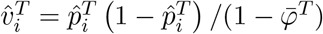, where 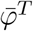 is the mean kinship estimated with `popkin`. Each causal locus is multiplied by -1 with probability 0.5. Alternatively, under the *random coefficients* (RC) model, initial causal coefficients are drawn independently from *β*_*i*_ ∼ Normal(0, 1). For both models, the initial genetic variance is 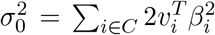, replacing 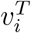 with 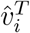 for unknown 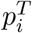 (so 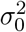 is an unbiased estimate), so we multiply every initial *β*_*i*_ by 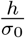 to have the desired heritability. Lastly, for known 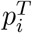, the intercept coefficient is 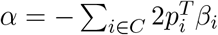. When 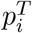 are unknown, 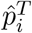 should not replace 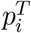 since that distorts the trait covariance (for the same reason the standard kinship estimator in Eq. (5) is biased), which is avoided with

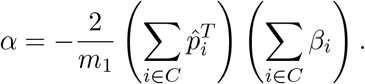

Simulations optionally included multiple environment group effects, similarly to previous models [40, 54], as follows. Each independent environment *i* has predefined groups, and each group *g* has random coefficients drawn independent from *η*_*gi*_ ∼ Normal(0, 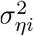) where 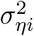 is a specified variance proportion for environment *i*. **Z** has individuals along columns and environment-groups along rows, and it contains indicator variables: 1 if the individual belongs to the environment-group, 0 otherwise. We performed trait simulations with the following variance parameters (Table 2): *high heritability* used *h*^2^ = 0.8 and no environment effects; *low heritability* used *h*^2^ = 0.3 and no environment effects; lastly, *environment* used *h*^2^ = 0.3, 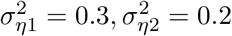 (total 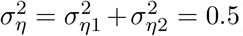). For real genotype datasets, the groups are the continental (environment 1) and fine-grained (environment 2) subpopulation labels given (see next subsection). For simulated genotypes, we created these labels by grouping by the index *j* (geographical coordinate) of each simulated individual, assigning group *g* = ceiling(*jk*_*i*_*/n*) where *k*_*i*_ is the number of groups in environment *i*, and we selected *k*_1_ = 5 and *k*_2_ = 25 to mimic the number of groups in each level of 1000 Genomes (Table 3).

**Table 2:** Variance parameters of trait simulations.

| Trait variance type | $h^2$ | $\sigma_{\eta}^2$ | $\sigma_{\epsilon}^2$ |
| --- | --- | --- | --- |
| High heritability | 0.8 | 0.0 | 0.2 |
| Low heritability | 0.3 | 0.0 | 0.7 |
| Environment | 0.3 | 0.5 | 0.2 |

**Table 3:**
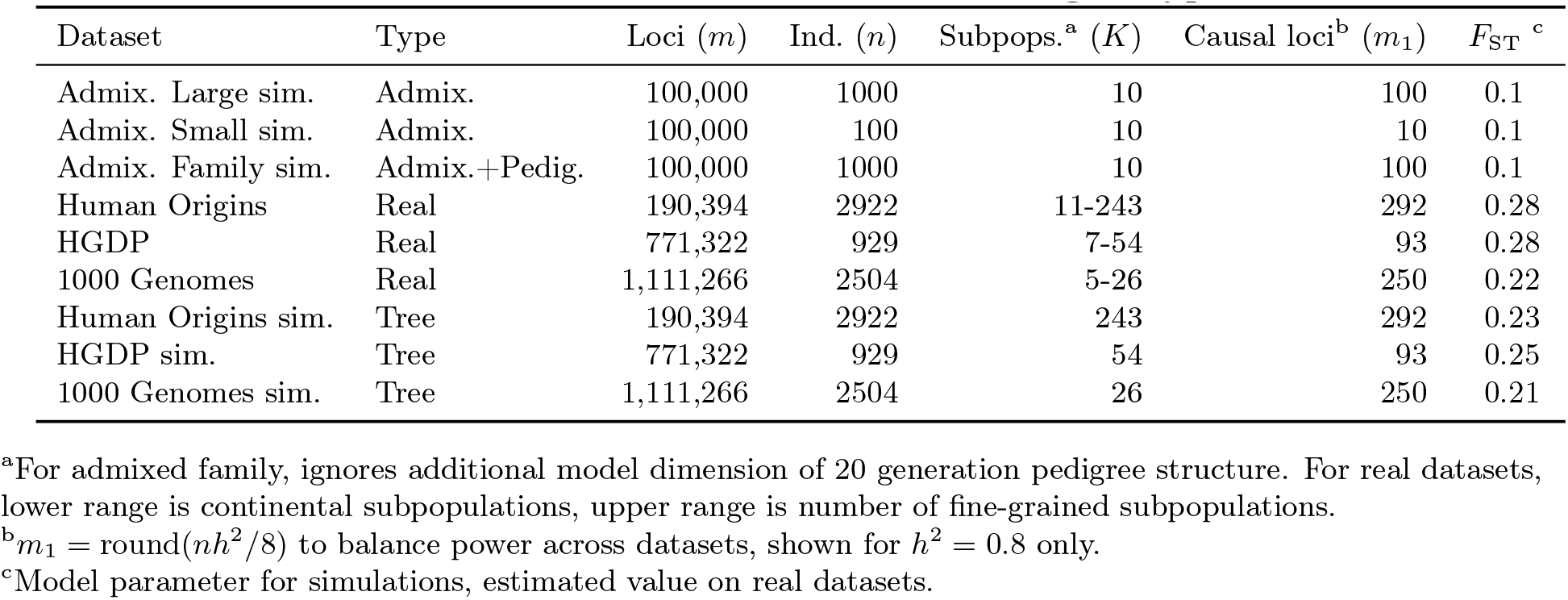
Features of simulated and real human genotype datasets.

### 2.3 Real human genotype datasets

The three datasets were processed as before [79] (summarized below), except with an additional filter so loci are in approximate linkage equilibrium and rare variants are removed. All processing was performed with `plink2` [96], and analysis was uniquely enabled by the R packages `BEDMatrix` [103] and `genio`. Each dataset groups individuals in a two-level hierarchy: continental and fine-grained subpopulations. Final dataset sizes are in Table 3.

We obtained the full (including non-public) Human Origins by contacting the authors and agreeing to their usage restrictions. The Pacific data [88] was obtained separately from the rest [86, 87], and datasets were merged using the intersection of loci. We removed ancient individuals, and individuals from singleton and non-native subpopulations. Non-autosomal loci were removed. Our analysis of both the whole-genome sequencing (WGS) version of HGDP [84] and the high-coverage NYGC version of 1000 Genomes [104] was restricted to autosomal biallelic SNP loci with filter “PASS”.

Since our evaluations assume uncorrelated loci, we filtered each real dataset with `plink2` using parameters “`--indep-pairwise 1000kb 0.3`”, which iteratively removes loci that have a greater than 0.3 squared correlation coefficient with another locus that is within 1000kb, stopping until no such loci remain. Since all real datasets have numerous rare variants, while PCA and LMM are not able to detect associations involving rare variants, we removed all loci with MAF < 0.01. Lastly, only HGDP had loci with over 10% missingness removed, as they were otherwise 17% of remaining loci (for Human Origins and 1000 Genomes they were under 1% of loci so they were not removed). Kinship matrix rank and eigenvalues were calculated from `popkin` kinship estimates. Eigenvalues were assigned p-values with `twstats` of the Eigensoft package [28], and kinship matrix rank was estimated as the largest number of consecutive eigenvalue from the start that all satisfy *p* < 0.01 (p-values did not increase monotonically). For the evaluation with close relatives removed, each dataset was filtered with `plink2` with option “`--king-cutoff`” with cutoff 0.02209709 (= 2^−11*/*2^) for removing up to 4th degree relatives using KING-robust [105], and MAF < 0.01 filter is reapplied (Table S1).

### 2.4 Evaluation of performance

All approaches are evaluated using two complementary metrics: SRMSD_*p*_ quantifies p-value uniformity, and AUC_PR_ measures causal locus classification performance and reflects power while ranking miscalibrated models fairly. These measures are more robust alternatives to previous measures from the literature (see Appendix B), and are implemented in `simtrait`.

P-values for continuous test statistics have a uniform distribution when the null hypothesis holds, a crucial assumption for type I error and FDR control [106, 107]. We use the Signed Root Mean Square Deviation (SRMSD_*p*_) to measure the difference between the observed null p-value quantiles and the expected uniform quantiles:

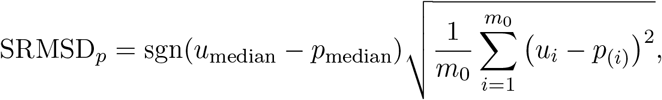

where *m*_0_ = *m* − *m*_1_ is the number of null (non-causal) loci, here *i* indexes null loci only, *p*_(*i*)_ is the *i*th ordered null p-value, *u*_*i*_ = (*i* − 0.5)*/m*_0_ is its expectation, *p*_median_ is the median observed null p-value, 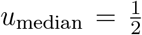 is its expectation, and sgn is the sign function (1 if *u*_median_ ≥ *p*_median_, -1 otherwise). Thus, SRMSD_*p*_ = 0 corresponds to calibrated p-values, SRMSD_*p*_ > 0 indicate anticonservative p-values, and SRMSD_*p*_ < 0 are conservative p-values. The maximum SRMSD_*p*_ is achieved when all p-values are zero (the limit of anti-conservative p-values), which for infinite loci approaches

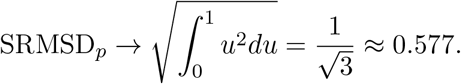

The same value with a negative sign occurs for all p-values of 1.

Precision and recall are standard performance measures for binary classifiers that do not require calibrated p-values [108]. Given the total numbers of true positives (TP), false positives (FP) and false negatives (FN) at some threshold or parameter *t*, precision and recall are

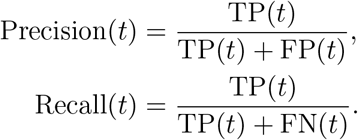

Precision and Recall trace a curve as *t* is varied, and the area under this curve is AUC_PR_. We use the R package `PRROC` to integrate the correct non-linear piecewise function when interpolating between points. A model obtains the maximum AUC_PR_ = 1 if there is a *t* that classifies all loci perfectly. In contrast, the worst models, which classify at random, have an expected precision (= AUC_PR_) equal to the overall proportion of causal loci: 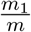.

## 3 Results

### 3.1 Overview of evaluations

We use three real genotype datasets and simulated genotypes from six population structure scenarios to cover various features of interest (Table 3). We introduce them in sets of three, as they appear in the rest of our results. Population kinship matrices, which combine population and family relatedness, are estimated without bias using `popkin` [52] (Fig. 1). The first set of three simulated genotypes are based on an admixture model with 10 ancestries (Fig. 1A) [35, 52, 109]. The “large” version (1000 individuals) illustrates asymptotic performance, while the “small” simulation (100 individuals) illustrates model overfitting. The “family” simulation has admixed founders and draws a 20-generation random pedigree with assortative mating, resulting in a complex joint family and ancestry structure in the last generation (Fig. 1B). The second set of three are the real human datasets representing global human diversity: Human Origins (Fig. 1D), HGDP (Fig. 1G), and 1000 Genomes (Fig. 1J), which are enriched for small minor allele frequencies even after MAF < 1% filter (Fig. 1C). Last are subpopulation tree simulations (Fig. 1F,I,L) fit to the kinship (Fig. 1E,H,K) and MAF (Fig. 1C) of each real human dataset, which by design do not have family structure.

**Figure 1:**
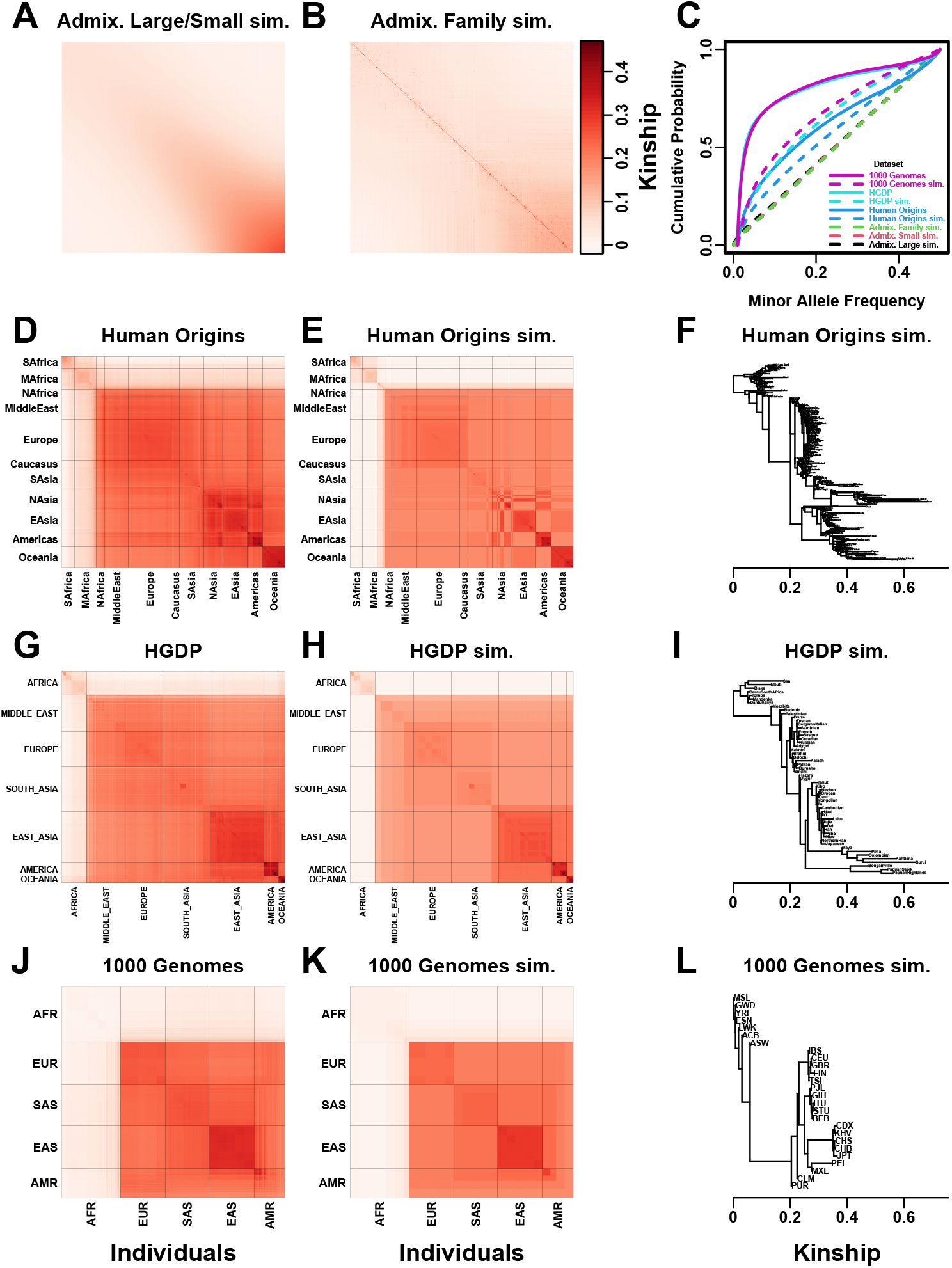
Population structures of simulated and real human genotype datasets. First two columns are population kinship matrices as heatmaps: individuals along x- and y-axis, kinship as color. Diagonal shows inbreeding values. **A**. Admixture scenario for both Large and Small simulations. **B**. Last generation of 20-generation admixed family, shows larger kinship values near diagonal corresponding to siblings, first cousins, etc. **C**. Minor allele frequency (MAF) distributions. Real datasets and subpopulation tree simulations had MAF ≥0.01 filter. **D**. Human Origins is an array dataset of a large diversity of global populations. **G**. Human Genome Diversity Panel (HGDP) is a WGS dataset from global native populations. **J**. 1000 Genomes Project is a WGS dataset of global cosmopolitan populations. **F**,**I**,**L**. Trees between subpopulations fit to real data. **E**,**H**,**K**. Simulations from trees fit to the real data recapitulate subpopulation structure.

All traits in this work are simulated. We repeated all evaluations on two additive quantitative trait models, *fixed effect sizes* (FES) and *random coefficients* (RC), which differ in how causal coefficients are constructed. The FES model captures the rough inverse relationship between coefficient and minor allele frequency that arises under strong negative and balancing selection and has been observed in numerous diseases and other traits [71, 89–91], so it is the focus of our results. The RC model draws coefficients independent of allele frequency, corresponding to neutral traits [71, 91], which results in a wider effect size distribution that reduces association power and effective polygenicity compared to FES.

We evaluate using two complementary measures: (1) SRMSD_*p*_ (p-value signed root mean square deviation) measures p-value calibration (closer to zero is better), and (2) AUC_PR_ (precision-recall area under the curve) measures causal locus classification performance (higher is better; Fig. 2). SRMSD_*p*_ is a more robust alternative to the common inflation factor *λ* and type I error control measures; there is a correspondence between *λ* and SRMSD_*p*_, with SRMSD_*p*_ > 0.01 giving *λ* > 1.06 (Fig. S1) and thus evidence of miscalibration close to the rule of thumb of *λ* > 1.05 [44]. There is also a monotonic correspondence between SRMSD_*p*_ and type I error rate (Fig. S2). AUC_PR_ has been used to evaluate association models [110], and reflects calibrated statistical power (Fig. S3) while being robust to miscalibrated models (Appendix B).

**Figure 2:**
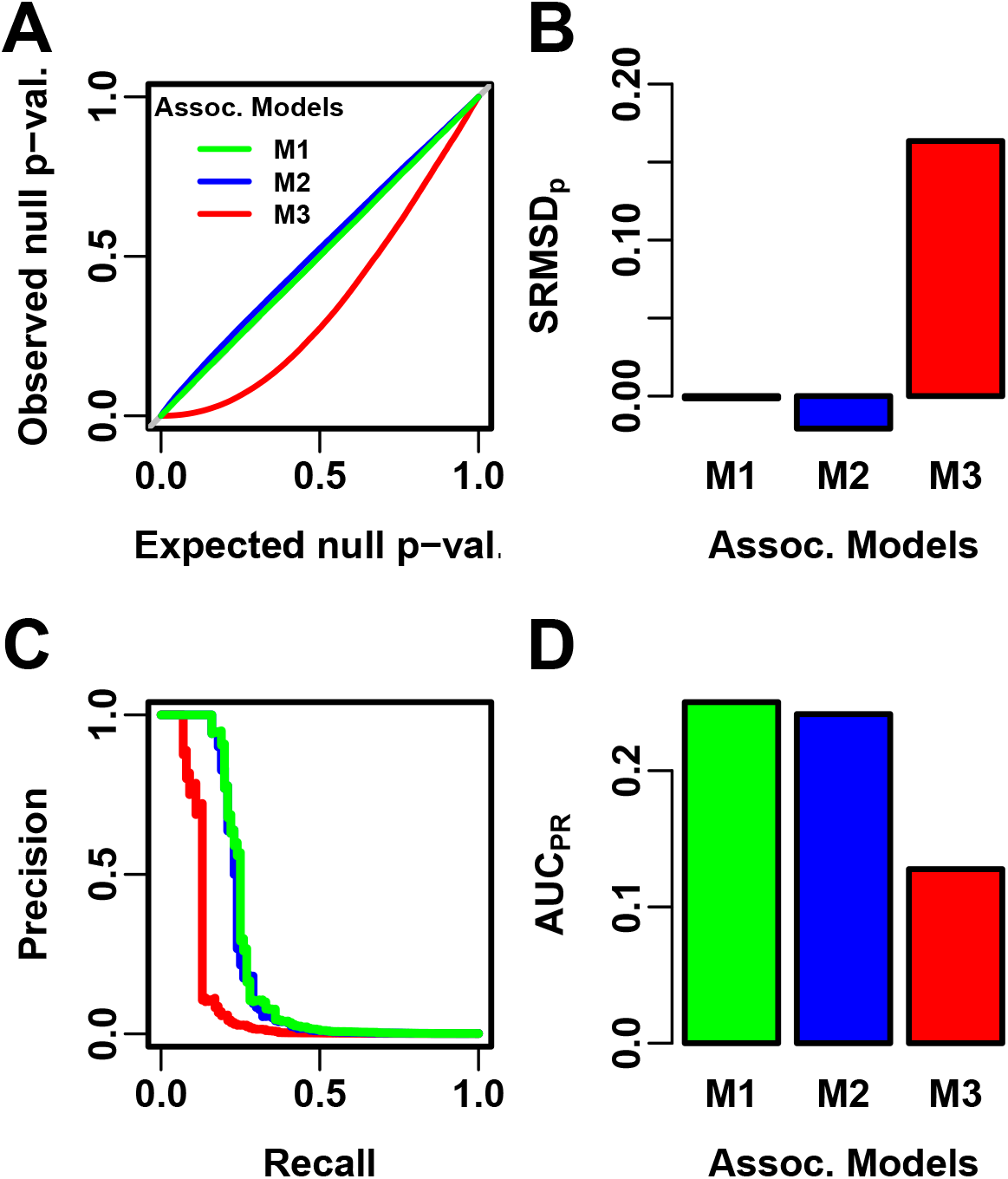
Illustration of evaluation measures. Three archetypal models illustrate our complementary measures: M1 is ideal, M2 overfits slightly, M3 is naive. **A**. QQ plot of p-values of “null” (non-causal) loci. M1 has desired uniform p-values, M2/M3 are miscalibrated. **B**. SRMSD_*p*_ (p-value Signed Root Mean Square Deviation) measures signed distance between observed and expected null p-values (closer to zero is better). **C**. Precision and Recall (PR) measure causal locus classification performance (higher is better). **D**. AUC_PR_ (Area Under the PR Curve) reflects power (higher is better).

Both PCA and LMM are evaluated in each replicate dataset including a number of PCs *r* between 0 and 90 as fixed covariates. In terms of p-value calibration, for PCA the best number of PCs *r* (minimizing mean |SRMSD_*p*_| over replicates) is typically large across all datasets (Table 4), although much smaller *r* values often performed as well (shown in following sections). Most cases have a mean |SRMSD_*p*_| < 0.01, whose p-values are effectively calibrated. However, PCA is often miscalibrated on the family simulation and real datasets (Table 4). In contrast, for LMM, *r* = 0 (no PCs) is always best, and is always calibrated. Comparing LMM with *r* = 0 to PCA with its best *r*, LMM always has significantly smaller |SRMSD_*p*_| than PCA or is statistically tied. For AUC_PR_ and PCA, the best *r* is always smaller than the best *r* for |SRMSD_*p*_|, so there is often a tradeoff between calibrated p-values versus classification performance. For LMM there is no tradeoff, as *r* = 0 often has the best mean AUC_PR_, and otherwise is not significantly different from the best *r*. Lastly, LMM with *r* = 0 always has significantly greater or statistically tied AUC_PR_ than PCA with its best *r*.

**Table 4:** Overview of PCA and LMM evaluations for high heritability simulations.

| Dataset | Metric | Trait<sup>a</sup> | LMM $r = 0$ vs best $r$ | | | PCA vs LMM $r = 0$ | | | |
| --- | --- | --- | --- | --- | --- | --- | --- | --- | --- |
| | | | Cal.<sup>b</sup> | Best $r^c$ | P-value<sup>d</sup> | Best $r^c$ | Cal.<sup>b</sup> | P-value<sup>d</sup> | Best model<sup>e</sup> |
| Admix. Large sim. | \|SRMSD<sub>p</sub>\| | FES | True | 0 | 1 | 12 | True | 0.036 | Tie |
| Admix. Small sim. | \|SRMSD<sub>p</sub>\| | FES | True | 0 | 1 | 4 | True | 0.055 | Tie |
| Admix. Family sim. | \|SRMSD<sub>p</sub>\| | FES | True | 0 | 1 | 90 | False | 3.9e-10* | LMM |
| Human Origins | \|SRMSD<sub>p</sub>\| | FES | True | 0 | 1 | 89 | False | 3.9e-10* | LMM |
| HGDP | \|SRMSD<sub>p</sub>\| | FES | True | 0 | 1 | 87 | True | 4.4e-10* | LMM |
| 1000 Genomes | \|SRMSD<sub>p</sub>\| | FES | True | 0 | 1 | 90 | False | 3.9e-10* | LMM |
| Human Origins sim. | \|SRMSD<sub>p</sub>\| | FES | True | 0 | 1 | 88 | True | 0.017 | Tie |
| HGDP sim. | \|SRMSD<sub>p</sub>\| | FES | True | 0 | 1 | 47 | True | 0.046 | Tie |
| 1000 Genomes sim. | \|SRMSD<sub>p</sub>\| | FES | True | 0 | 1 | 78 | True | 9.6e-10* | LMM |
| Admix. Large sim. | \|SRMSD<sub>p</sub>\| | RC | True | 0 | 1 | 26 | True | 0.11 | Tie |
| Admix. Small sim. | \|SRMSD<sub>p</sub>\| | RC | True | 0 | 1 | 4 | True | 0.00097 | Tie |
| Admix. Family sim. | \|SRMSD<sub>p</sub>\| | RC | True | 0 | 1 | 90 | False | 3.9e-10* | LMM |
| Human Origins | \|SRMSD<sub>p</sub>\| | RC | True | 0 | 1 | 90 | True | 0.00065 | Tie |
| HGDP | \|SRMSD<sub>p</sub>\| | RC | True | 0 | 1 | 37 | True | 1.5e-05* | LMM |
| 1000 Genomes | \|SRMSD<sub>p</sub>\| | RC | True | 0 | 1 | 76 | True | 3.9e-10* | LMM |
| Human Origins sim. | \|SRMSD<sub>p</sub>\| | RC | True | 0 | 1 | 85 | True | 0.14 | Tie |
| HGDP sim. | \|SRMSD<sub>p</sub>\| | RC | True | 0 | 1 | 44 | True | 8.8e-07* | LMM |
| 1000 Genomes sim. | \|SRMSD<sub>p</sub>\| | RC | True | 0 | 1 | 90 | True | 3.9e-10* | LMM |
| Admix. Large sim. | AUC<sub>PR</sub> | FES |  | 0 | 1 | 3 |  | 5.9e-06* | LMM |
| Admix. Small sim. | AUC<sub>PR</sub> | FES |  | 0 | 1 | 2 |  | 0.025 | Tie |
| Admix. Family sim. | AUC<sub>PR</sub> | FES |  | 1 | 0.35 | 22 |  | 3.9e-10* | LMM |
| Human Origins | AUC<sub>PR</sub> | FES |  | 0 | 1 | 34 |  | 3.9e-10* | LMM |
| HGDP | AUC<sub>PR</sub> | FES |  | 1 | 0.33 | 16 |  | 4.4e-10* | LMM |
| 1000 Genomes | AUC<sub>PR</sub> | FES |  | 1 | 0.11 | 8 |  | 3.9e-10* | LMM |
| Human Origins sim. | AUC<sub>PR</sub> | FES |  | 0 | 1 | 36 |  | 3.9e-10* | LMM |
| HGDP sim. | AUC<sub>PR</sub> | FES |  | 0 | 1 | 17 |  | 1.7e-05* | LMM |
| 1000 Genomes sim. | AUC<sub>PR</sub> | FES |  | 0 | 1 | 10 |  | 5e-10* | LMM |
| Admix. Large sim. | AUC<sub>PR</sub> | RC |  | 0 | 1 | 3 |  | 1.4e-05* | LMM |
| Admix. Small sim. | AUC<sub>PR</sub> | RC |  | 0 | 1 | 1 |  | 0.095 | Tie |
| Admix. Family sim. | AUC<sub>PR</sub> | RC |  | 0 | 1 | 34 |  | 3.9e-10* | LMM |
| Human Origins | AUC<sub>PR</sub> | RC |  | 3 | 0.4 | 36 |  | 9.6e-10* | LMM |
| HGDP | AUC<sub>PR</sub> | RC |  | 4 | 0.21 | 16 |  | 0.013 | Tie |
| 1000 Genomes | AUC<sub>PR</sub> | RC |  | 5 | 0.004 | 9 |  | 0.00043 | Tie |
| Human Origins sim. | AUC<sub>PR</sub> | RC |  | 0 | 1 | 37 |  | 4.1e-10* | LMM |
| HGDP sim. | AUC<sub>PR</sub> | RC |  | 3 | 0.087 | 17 |  | 0.0014 | Tie |
| 1000 Genomes sim. | AUC<sub>PR</sub> | RC |  | 3 | 0.37 | 10 |  | 8.5e-10* | LMM |

<sup>a</sup>FES: Fixed Effect Sizes, RC: Random Coefficients.<sup>b</sup>Calibrated: whether mean |SRMSD<sub>p</sub>| < 0.01.<sup>c</sup>Value of $r$ (number of PCs) with minimum mean |SRMSD<sub>p</sub>| or maximum mean AUC<sub>PR</sub>.<sup>d</sup>Wilcoxon paired 1-tailed test of distributions (|SRMSD<sub>p</sub>| or AUC<sub>PR</sub>) between models in header. Asterisk marks significant value using Bonferroni threshold ( $p < \alpha/n_{\text{tests}}$ with $\alpha = 0.01$ and $n_{\text{tests}} = 72$ is the number of tests in this table).<sup>e</sup>Tie if no significant difference using Bonferroni threshold.

### 3.2 Evaluations in admixture simulations

Now we look more closely at results per dataset. The complete SRMSD_*p*_ and AUC_PR_ distributions for the admixture simulations and FES traits are in Fig. 3. RC traits gave qualitatively similar results (Fig. S4).

**Figure 3:**
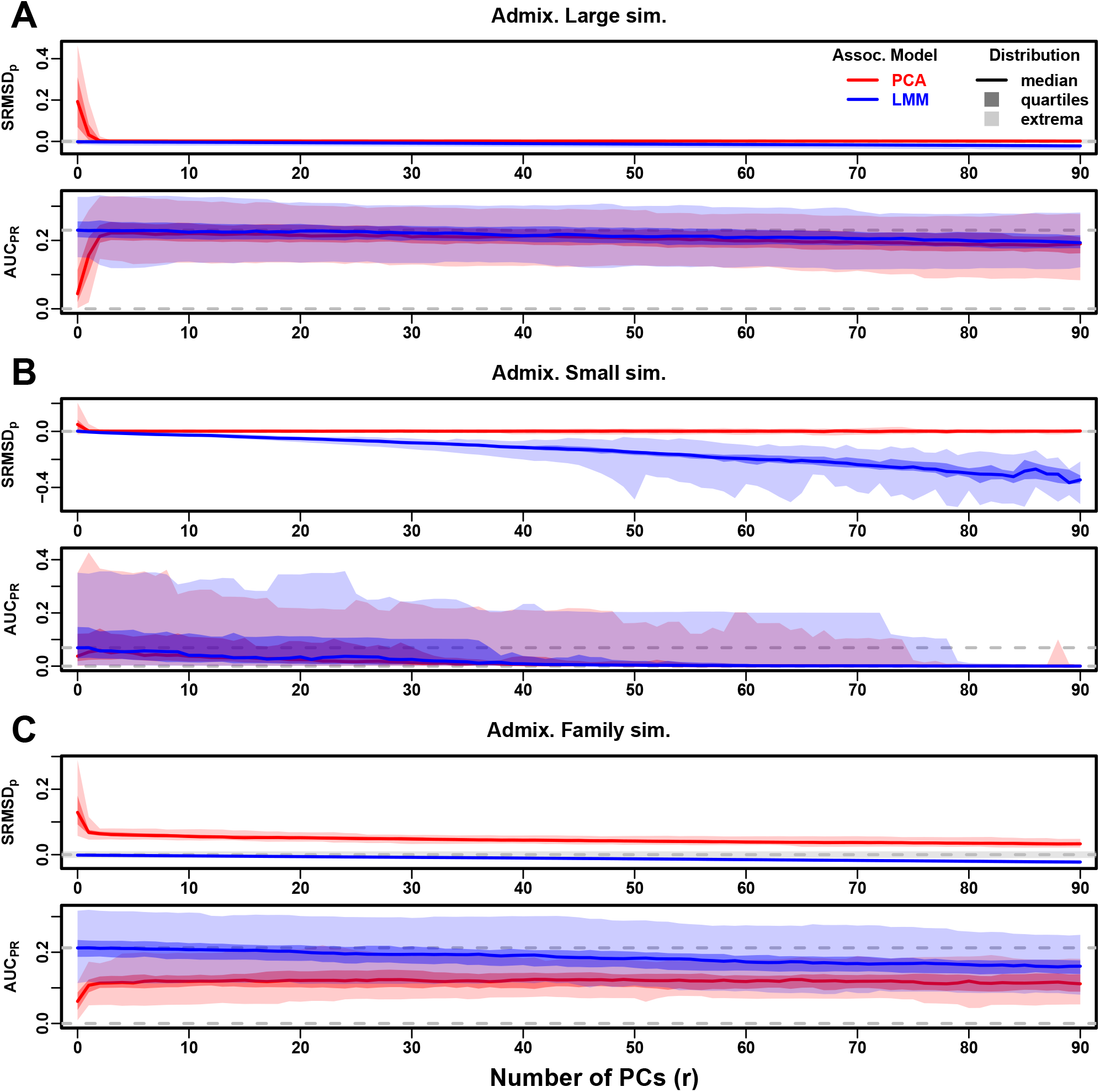
Evaluations in admixture simulations. Traits simulated from FES model with high heritability. PCA and LMM models have varying number of PCs (*r* ∈ {0, …, 90} on x-axis), with the distributions (y-axis) of SRMSD_*p*_ (top subpanel) and AUC_PR_ (bottom subpanel) for 50 replicates. Best performance is zero SRMSD_*p*_ and large AUC_PR_. Zero and maximum median AUC_PR_ values are marked with horizontal gray dashed lines, and |SRMSD_*p*_| < 0.01 is marked with a light gray area. LMM performs best with *r* = 0, PCA with various *r*. **A**. Large simulation (*n* = 1, 000 individuals). **B**. Small simulation (*n* = 100) shows overfitting for large *r*. **C**. Family simulation (*n* = 1, 000) has admixed founders and large numbers of close relatives from a realistic random 20-generation pedigree. PCA performs poorly compared to LMM: SRMSD_*p*_ > 0 for all *r* and large AUC_PR_ gap.

In the large admixture simulation, the SRMSD_*p*_ of PCA is largest when *r* = 0 (no PCs) and decreases rapidly to near zero at *r* = 3, where it stays for up to *r* = 90 (Fig. 3A). Thus, PCA has calibrated p-values for *r* ≥ 3, smaller than the theoretical optimum for this simulation of *r* = *K* − 1 = 9. In contrast, the SRMSD_*p*_ for LMM starts near zero for *r* = 0, but becomes negative as *r* increases (p-values are conservative). The AUC_PR_ distribution of PCA is similarly worst at *r* = 0, increases rapidly and peaks at *r* = 3, then decreases slowly for *r* > 3, while the AUC_PR_ distribution for LMM starts near its maximum at *r* = 0 and decreases with *r*. Although the AUC_PR_ distributions for LMM and PCA overlap considerably at each *r*, LMM with *r* = 0 has significantly greater AUC_PR_ values than PCA with *r* = 3 (Table 4). However, qualitatively PCA performs nearly as well as LMM in this simulation.

The observed robustness to large *r* led us to consider smaller sample sizes. A model with large numbers of parameters *r* should overfit more as *r* approaches the sample size *n*. Rather than increase *r* beyond 90, we reduce individuals to *n* = 100, which is small for typical association studies but may occur in studies of rare diseases, pilot studies, or other constraints. To compensate for the loss of power due to reducing *n*, we also reduce the number of causal loci (see Materials and Methods), which increases per-locus effect sizes. We found a large decrease in performance for both models as *r* increases, and best performance for *r* = 1 for PCA and *r* = 0 for LMM (Fig. 3B). Remarkably, LMM attains much larger negative SRMSD_*p*_ values than in our other evaluations. LMM with *r* = 0 is significantly better than PCA (*r* = 1 to 4) in both measures (Table 4), but qualitatively the difference is negligible.

The family simulation adds a 20-generation random family to our large admixture simulation. Only the last generation is studied for association, which contains numerous siblings, first cousins, etc., with the initial admixture structure preserved by geographically biased mating. Our evaluation reveals a sizable gap in both measures between LMM and PCA across all *r* (Fig. 3C). LMM again performs best with *r* = 0 and achieves mean |SRMSD_*p*_| < 0.01. However, PCA does not achieve mean |SRMSD_*p*_| < 0.01 at any *r*, and its best mean AUC_PR_ is considerably worse than that of LMM. Thus, LMM is conclusively superior to PCA, and the only calibrated model, when there is family structure.

### 3.3 Evaluations in real human genotype datasets

Next we repeat our evaluations with real human genotype data, which differs from our simulations in allele frequency distributions and more complex population structures with greater *F*_ST_, numerous correlated subpopulations, and potential cryptic family relatedness.

Human Origins has the greatest number and diversity of subpopulations. The SRMSD_*p*_ and AUC_PR_ distributions in this dataset and FES traits (Fig. 4A) most resemble those from the family simulation (Fig. 3C). In particular, while LMM with *r* = 0 performed optimally (both measures) and satisfies mean |SRMSD_*p*_| < 0.01, PCA maintained SRMSD_*p*_ > 0.01 for all *r* and its AUC_PR_ were all considerably smaller than the best AUC_PR_ of LMM.

**Figure 4:**
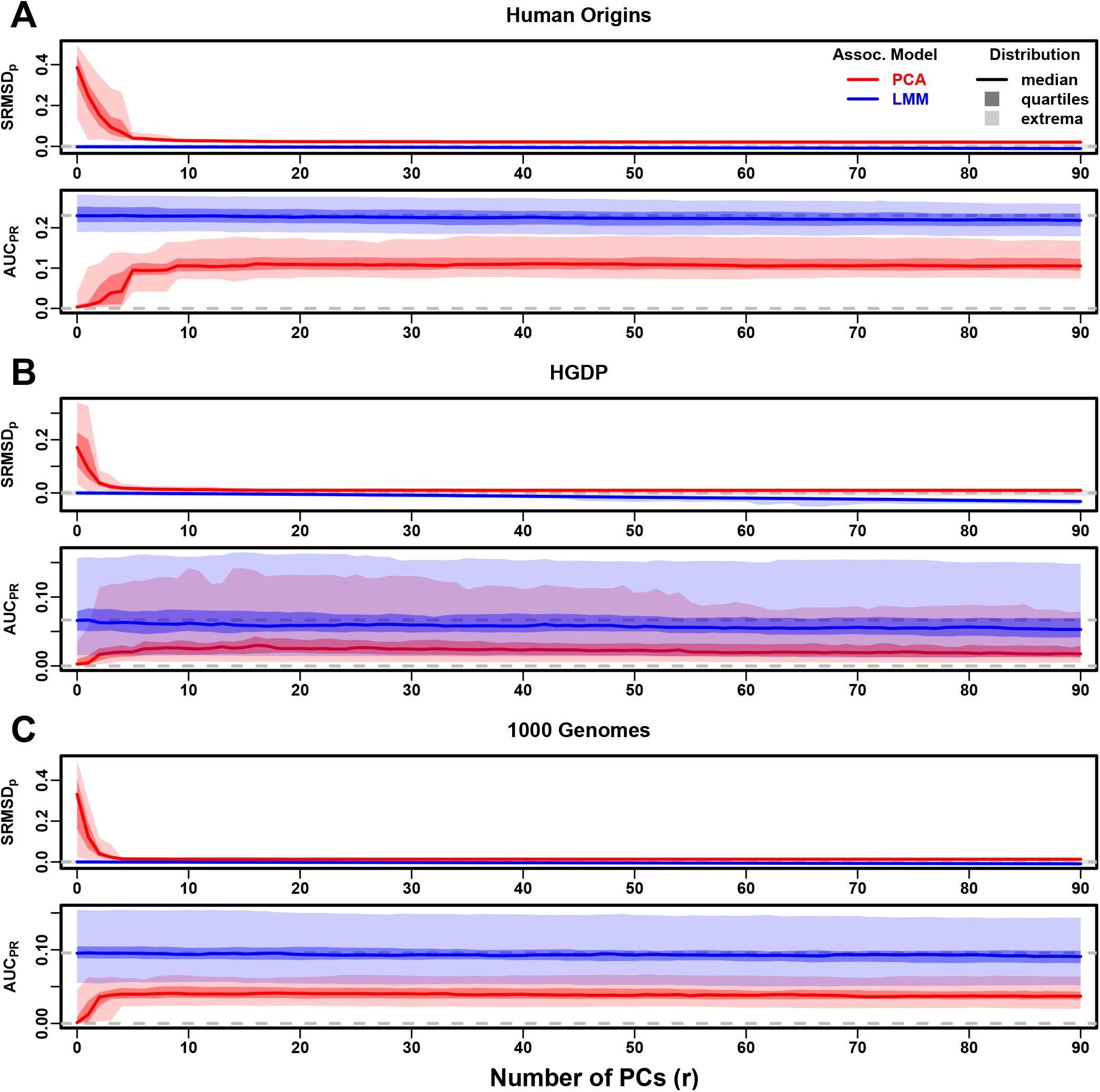
Evaluations in real human genotype datasets. Traits simulated from FES model with high heritability. Same setup as Fig. 3, see that for details. These datasets strongly favor LMM with no PCs over PCA, with distributions that most resemble the family simulation. **A**. Human Origins. **B**. Human Genome Diversity Panel (HGDP). **C**. 1000 Genomes Project.

HGDP has the fewest individuals among real datasets, but compared to Human Origins contains more loci and low-frequency variants. Performance (Fig. 4B) again most resembled the family simulations. In particular, LMM with *r* = 0 achieves mean |SRMSD_*p*_| < 0.01 (p-values are calibrated), while PCA does not, and there is a sizable AUC_PR_ gap between LMM and PCA. Maximum AUC_PR_ values were lowest in HGDP compared to the two other real datasets.

1000 Genomes has the fewest subpopulations but largest number of individuals per subpopulation. Thus, although this dataset has the simplest subpopulation structure among the real datasets, we find SRMSD_*p*_ and AUC_PR_ distributions (Fig. 4C) that again most resemble our earlier family simulation, with mean |SRMSD_*p*_| < 0.01 for LMM only and large AUC_PR_ gaps between LMM and PCA.

Our results are qualitatively different for RC traits, which had smaller AUC_PR_ gaps between LMM and PCA (Fig. S5). Maximum AUC_PR_ were smaller in RC compared to FES in Human Origins and 1000 Genomes, suggesting lower power for RC traits across association models. Nevertheless, LMM with *r* = 0 was significantly better than PCA for all measures in the real datasets and RC traits (Table 4).

### 3.4 Evaluations in subpopulation tree simulations fit to human data

To better understand which features of the real datasets lead to the large differences in performance between LMM and PCA, we carried out subpopulation tree simulations. Human subpopulations are related roughly by trees, which induce the strongest correlations, so we fit trees to each real dataset and tested if data simulated from these complex tree structures could recapitulate our previous results (Fig. 1). These tree simulations also feature non-uniform ancestral allele frequency distributions, which recapitulated some of the skew for smaller minor allele frequencies of the real datasets (Fig. 1C). The SRMSD_*p*_ and AUC_PR_ distributions for these tree simulations (Fig. 5) resembled our admixture simulation more than either the family simulation (Fig. 3) or real data results (Fig. 4). Both LMM with *r* = 0 and PCA (various *r*) achieve mean |SRMSD_*p*_| < 0.01 (Table 4). The AUC_PR_ distributions of both LMM and PCA track closely as *r* is varied, although there is a small gap resulting in LMM (*r* = 0) besting PCA in all three simulations. The results are qualitatively similar for RC traits (Fig. S6 and Table 4). Overall, these subpopulation tree simulations do not recapitulate the large LMM advantage over PCA observed on the real data.

**Figure 5:**
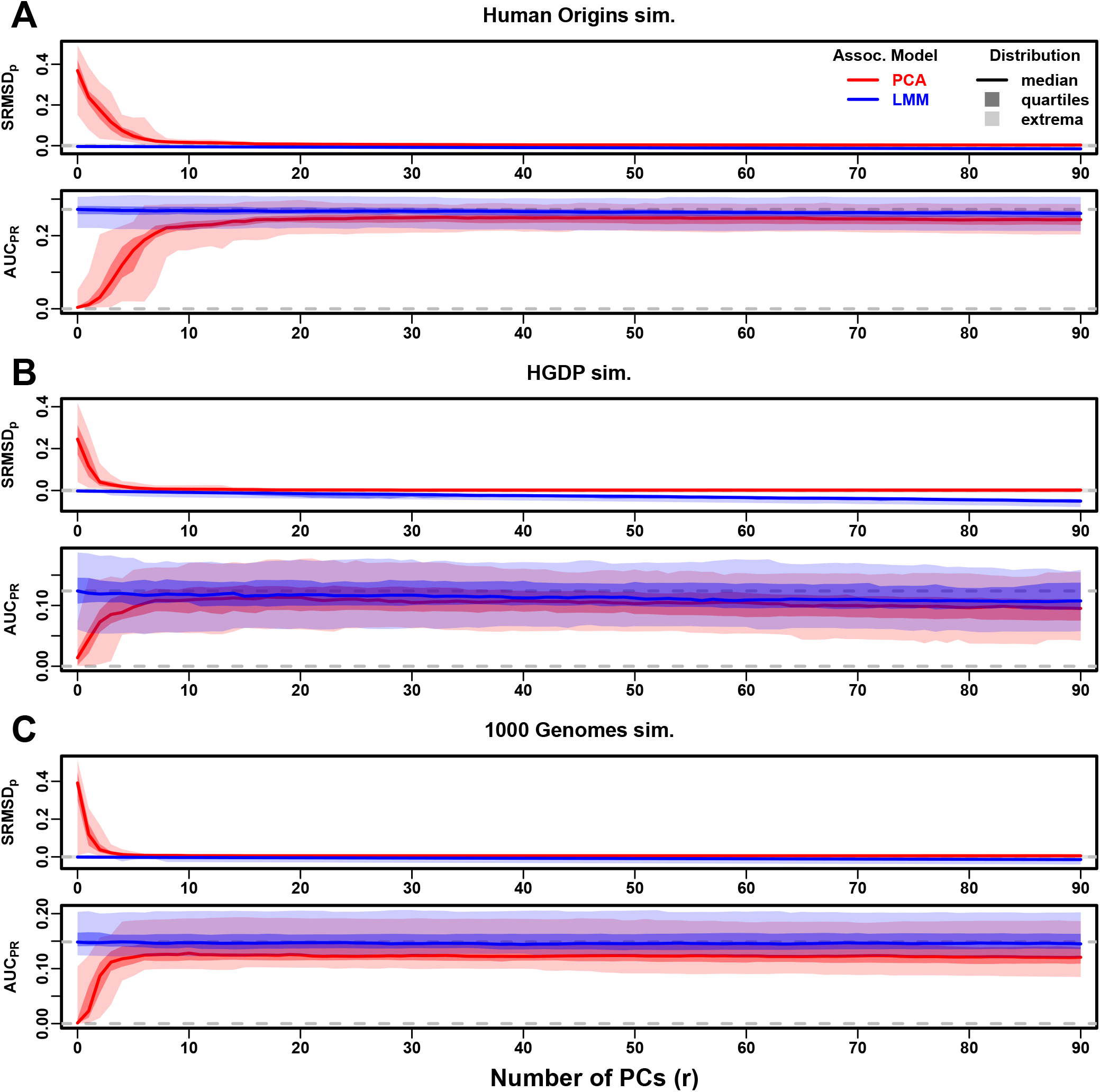
Evaluations in subpopulation tree simulations fit to human data. Traits simulated from FES model with high heritability. Same setup as Fig. 3, see that for details. These tree simulations, which exclude family structure by design, do not explain the large gaps in LMM-PCA performance observed in the real data. **A**. Human Origins tree simulation. **B**. Human Genome Diversity Panel (HGDP) tree simulation. **C**. 1000 Genomes Project tree simulation.

### 3.5 Numerous distant relatives explain poor PCA performance in real data

In principle, PCA performance should be determined by the dimension of relatedness, or kinship matrix rank, since PCA is a low-dimensional model whereas LMM can model high-dimensional relatedness without overfitting. We used the Tracy-Widom test [28] with *p* < 0.01 to estimate kinship matrix rank as the number of significant PCs (Fig. S7A). The true rank of our simulations is slightly underestimated (Table 3), but we confirm that the family simulation has the greatest rank, and real datasets have greater estimates than their respective subpopulation tree simulations, which confirms our hypothesis to some extent. However, estimated ranks do not separate real datasets from tree simulations, as required to predict the observed PCA performance. Moreover, the HGDP and 1000 Genomes rank estimates are 45 and 61, respectively, yet PCA performed poorly for all *r* ≤ 90 numbers of PCs (Fig. 4). The top eigenvalue explained a proportion of variance proportional to *F*_ST_ (Table 3), but the rest of the top 10 eigenvalues show no clear differences between datasets, except the small simulation had larger variances explained per eigenvalue (expected since it has fewer eigenvalues; Fig. S7C). Comparing cumulative variance explained versus rank fraction across all eigenvalues, all datasets increase from their starting point almost linearly until they reach 1, except the family simulation has much greater variance explained by mid-rank eigenvalues (Fig. S7B). We also calculated the number of PCs that are significantly associated with the trait, and observed similar results, namely that while the family simulation has more significant PCs than the nonfamily admixture simulations, the real datasets and their tree simulated counterparts have similar numbers of significant PCs (Fig. S8). Overall, there is no separation between real datasets (where PCA performed poorly) and subpopulation tree simulations (where PCA performed relatively well) in terms of their eigenvalues or kinship matrix rank estimates.

Local kinship, which is recent relatedness due to family structure excluding population structure, is the presumed cause of the LMM to PCA performance gap observed in real datasets but not their subpopulation tree simulation counterparts. Instead of inferring local kinship through increased kinship matrix rank, as attempted in the last paragraph, now we measure it directly using the KING-robust estimator [105]. We observe more large local kinship in the real datasets and the family simulation compared to the other simulations (Fig. 6). However, for real data this distribution depends on the subpopulation structure, since locally related pairs are most likely in the same subpopulation. Therefore, the only comparable curve to each real dataset is their corresponding subpopulation tree simulation, which matches subpopulation structure. In all real datasets we identified highly related individual pairs with kinship above the 4th degree relative threshold of 0.022 [105, 111]. However, these highly related pairs are vastly outnumbered by more distant pairs with evident non-zero local kinship as compared to the extreme tree simulation values.

**Figure 6:**
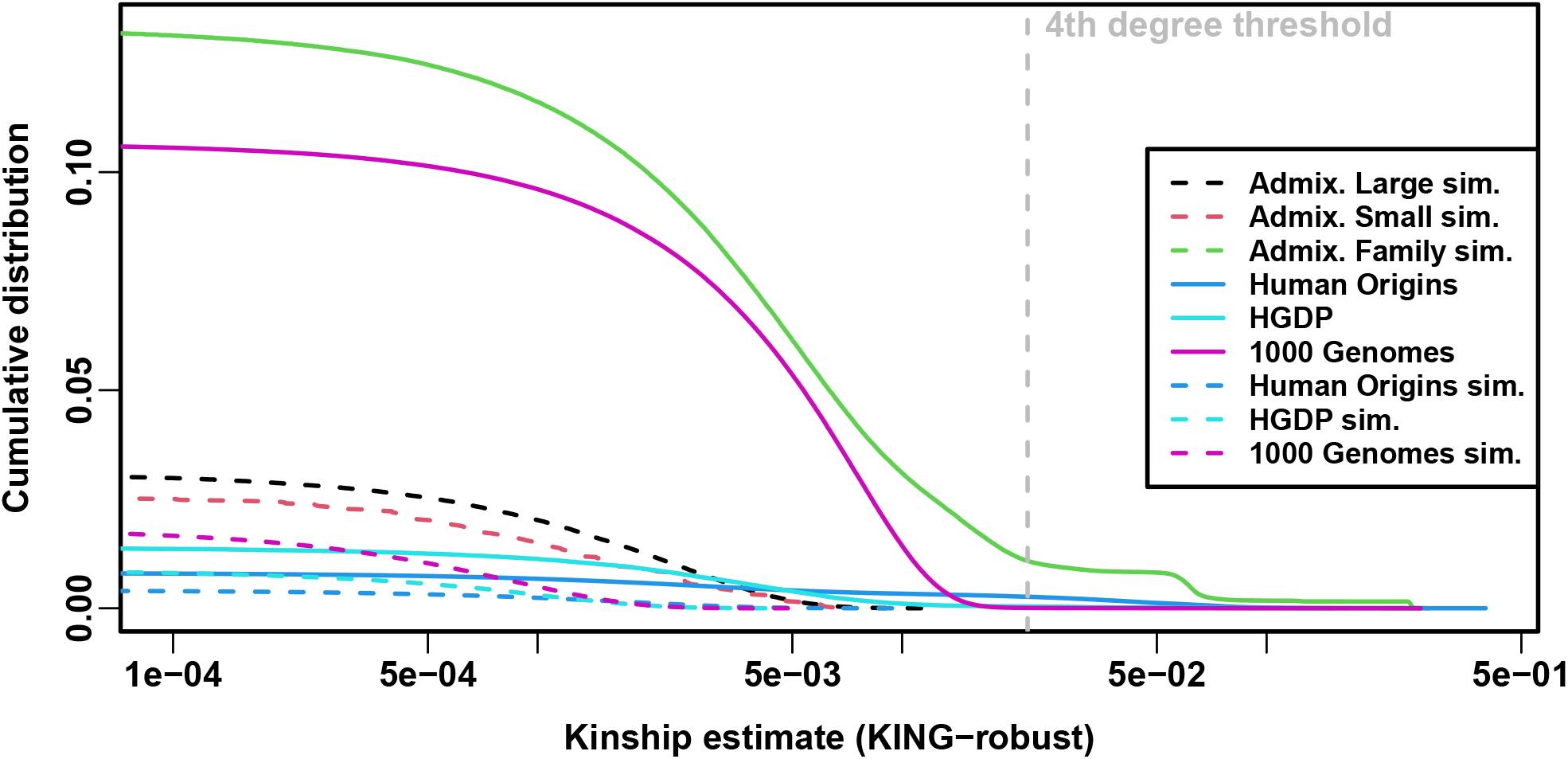
Local kinship distributions. Curves are complementary cumulative distribution of lower triangular kinship matrix (self kinship excluded) from KING-robust estimator. Note log x-axis; negative estimates are counted but not shown. Most values are below 4th degree relative threshold. Each real dataset has a greater cumulative than its subpopulation tree simulations.

To try to improve PCA performance, we followed the standard practice of removing 4th degree relatives, which reduced sample sizes between 5% and 10% (Table S1). Only *r* = 0 for LMM and *r* = 20 for PCA were tested, as these performed well in our earlier evaluation, and only FES traits were tested because they previously displayed the large PCA-LMM performance gap. LMM significantly outperforms PCA in all these cases (Wilcoxon paired 1-tailed *p* < 0.01; Fig. 7). Notably, PCA still had miscalibrated p-values two of the three real datasets (|SRMSD_*p*_| > 0.01), the only marginally calibrated case being HGDP which is also the smallest of these datasets. Otherwise, AUC_PR_ and SRMSD_*p*_ ranges were similar here as in our earlier evaluation. Therefore, the removal of the small number of highly related individual pairs had a negligible effect in PCA performance, so the larger number of more distantly related pairs explain the poor PCA performance in the real datasets.

**Figure 7:**
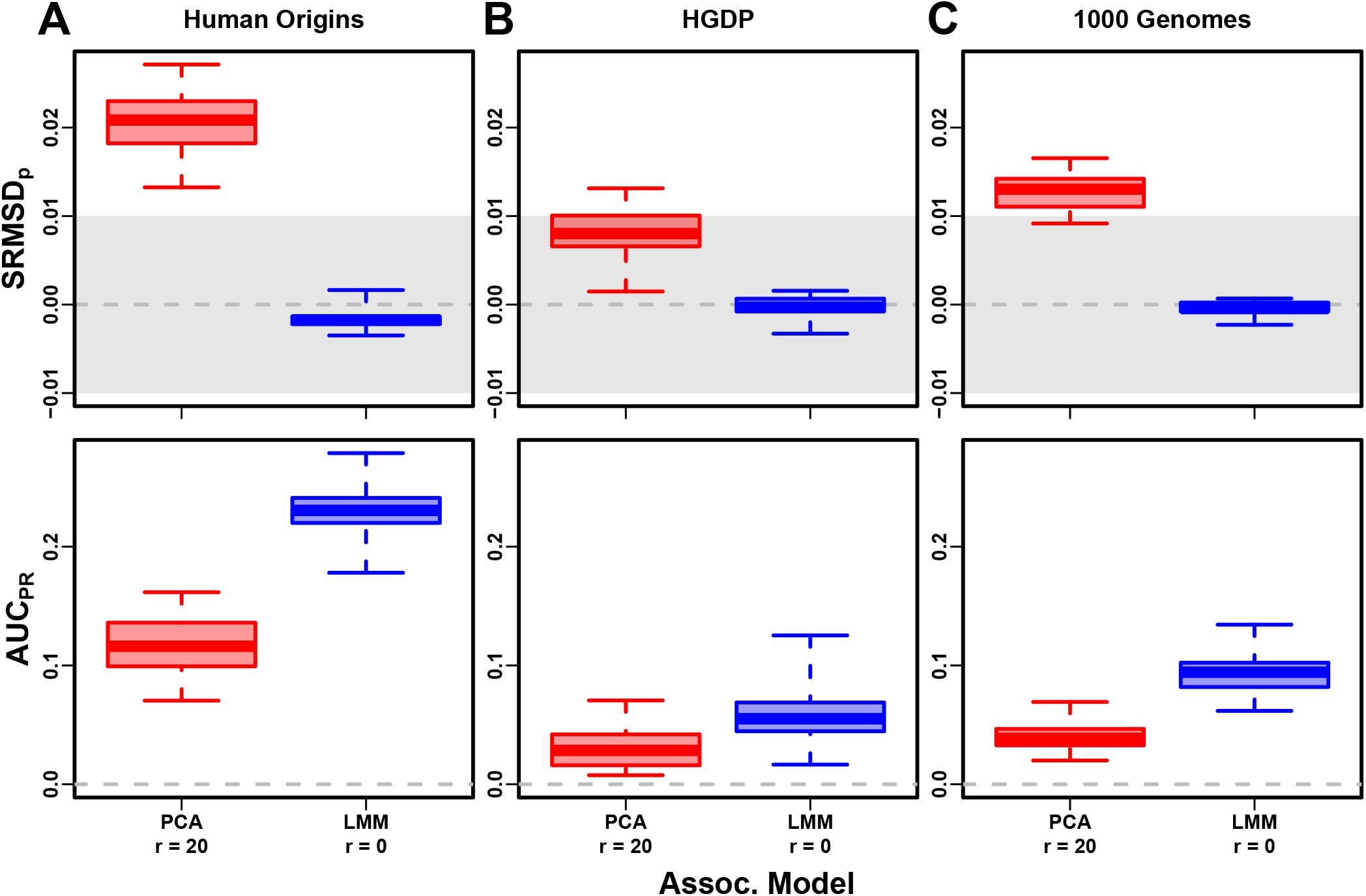
Evaluation in real datasets excluding 4th degree relatives. Traits simulated from FES model with high heritability. Each dataset is a column, rows are measures. First row has |SRMSD_*p*_| < 0.01 band marked as gray area.

### 3.6 Low heritability and environment simulations

Our main evaluations were repeated with traits simulated under a lower heritability value of *h*^2^ = 0.3. We reduced the number of causal loci in response to this change in heritability, to result in equal average effect size per locus compared to the previous high heritability evaluations (see Materials and Methods). Despite that, these low heritability evaluations measured lower AUC_PR_ values than their high heritability counterparts (Figs. S9 to S13). The gap between LMM and PCA was reduced in these evaluations, but the main conclusion of the high heritability evaluation holds for low heritability as well, namely that LMM with *r* = 0 significantly outperforms or ties LMM with *r* > 0 and PCA in all cases (Table S2).

Lastly, we simulated traits with both low heritability and large environment effects determined by geography and subpopulation labels, so they are strongly correlated to the low-dimensional population structure (Table 2). For that reason, PCs may be expected to perform better in this setting (in either PCA or LMM). However, we find that both PCA and LMM (even without PCs) increase their AUC_PR_ values compared to the low-heritability evaluations (Fig. S14; Fig. 8 also shows representative numbers of PCs, which performed optimally or nearly so in individual simulations shown in Figs. S15 to S18). P-value calibration is comparable with or without environment effects, for LMM for all *r* and for PCA once *r* is large enough (Fig. S14). These simulations are the only where we occasionally observed for both metrics a significant, though small, advantage of LMM with PCs versus LMM without PCs (Table S3). Additionally, on RC traits only, PCA significantly outperforms LMM in the three real human datasets (Table S3), the only cases in all of our evaluations where this is observed. For comparison, we also evaluate an “oracle” LMM without PCs but with the finest group labels, the same used to simulate environment, as fixed categorical covariates (“LMM lab.”), and see much larger AUC_PR_ values than either LMM with PCs or PCA (Figs. 8 and S15 to S18 and Table S3). However, LMM with labels is often more poorly calibrated than LMM or PCA without labels, which may be since these numerous labels are inappropriately modeled as fixed rather than random effects. Overall, we find that association studies with correlated environment and genetic effects remain a challenge for PCA and LMM, that addition of PCs to an LMM improves performance only marginally, and that if the environment effect is driven by geography or ethnicity then use of those labels greatly improves performance compared to using PCs.

**Figure 8:**
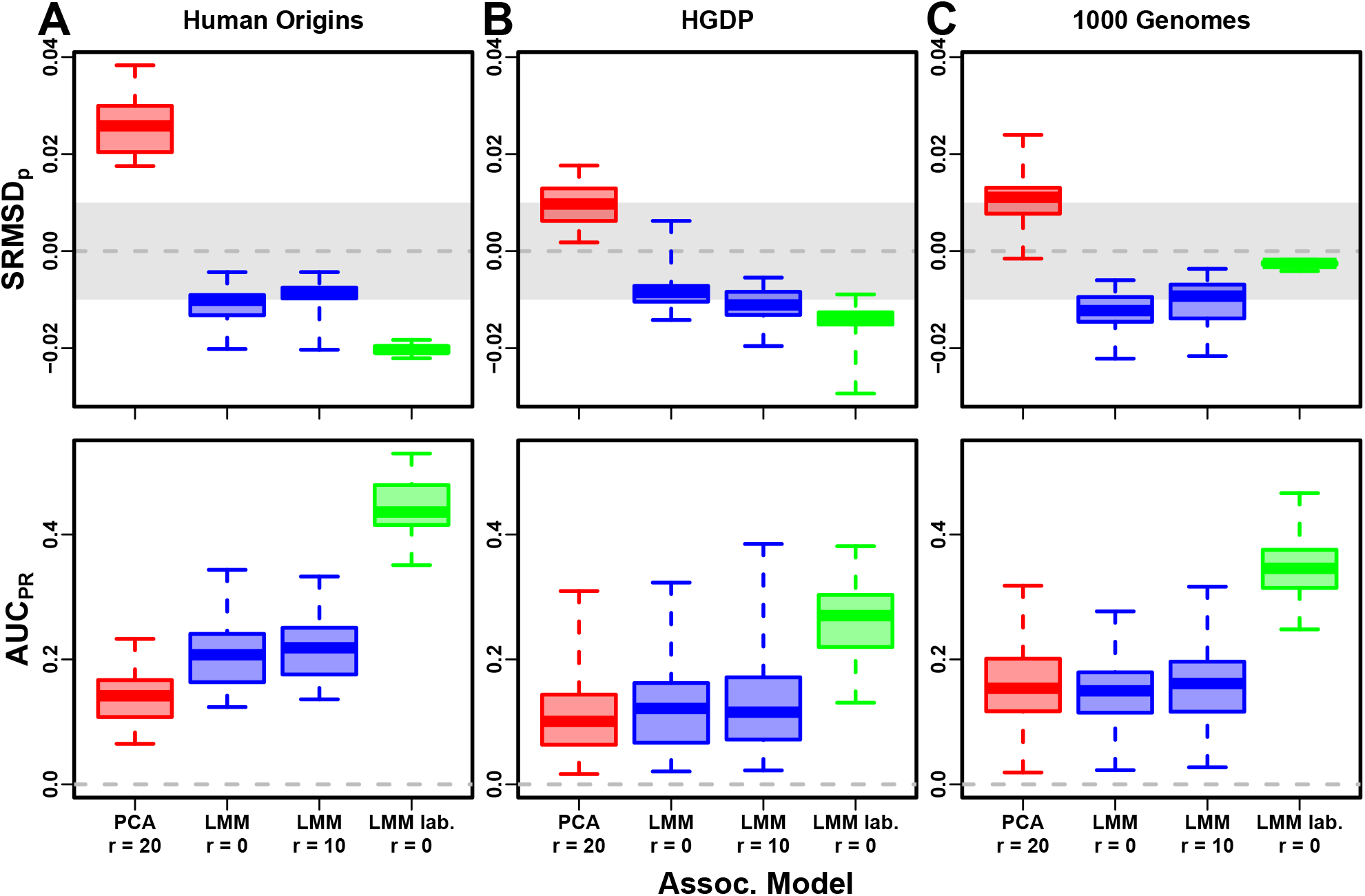
Evaluation in real datasets excluding 4th degree relatives, FES traits, environment. Traits simulated with environment effects, otherwise the same as Fig. 7. “LMM lab.” includes as fixed effects true groups from which environment was simulated.

## 4 Discussion

Our evaluations conclusively determined that LMM without PCs performs better than PCA (for any number of PCs) across all scenarios without environment effects, including all real and simulated genotypes and two trait simulation models. Although the addition of a few PCs to LMM does not greatly hurt its performance (except for small sample sizes), they generally did not improve it either (Tables S2 and 4), which agrees with previous observations [68, 70] but contradicts others [37, 44]. Our findings make sense since PCs are the eigenvectors of the same kinship matrix that parameterized random effects, so including both is redundant.

The presence of environment effects that are correlated to relatedness presents the only scenario where occasionally PCA and LMM with PCs outperform LMM without PCs (Table S3). It is commonly believed that PCs model such environment effects well [20, 39, 40]. However, we observe that LMM without PCs models environment effects nearly as well as with PCs (Fig. 8), consistent with previous findings [53, 54] and with environment inflating heritability estimates using LMM [112]. Moreover, modeling the true environment groups as fixed categorical effects always substantially improved AUC_PR_ compared to modeling them with PCs (Fig. 8 and Table S3). Modeling numerous environment groups as fixed effects does result in deflated p-values (Fig. 8 and Table S3), which we expect would be avoided by modeling them as random effects, a strategy we chose not to pursue here as it is both a circular evaluation (the true effects were drawn from that model) and out of scope. Overall, including PCs to model environment effects yields limited power gains if at all, even in an LMM, and is no replacement for more adequate modeling of environment whenever possible. Previous studies found that PCA was better calibrated than LMM for unusually differentiated markers [44, 55, 57], which as simulated were an artificial scenario not based on a population genetics model, and are otherwise believed to be unusual [58, 78]. Our evaluations on real human data, which contain such loci in relevant proportions if they exist, do not replicate that result. Family relatedness strongly favors LMM, an advantage that probably outweighs this potential PCA benefit in real data.

Relative to LMM, the behavior of PCA fell between two extremes. When PCA performed well, there was a small number of PCs with both calibrated p-values and AUC_PR_ near that of LMM without PCs. Conversely, PCA performed poorly when no number of PCs had either calibrated p-values or acceptably large AUC_PR_. There were no cases where high numbers of PCs optimized an acceptable AUC_PR_, or cases with miscalibrated p-values but high AUC_PR_. PCA performed well in the admixture simulations (without families, both trait models), real human genotypes with RC traits, and the subpopulation tree simulations (both trait models). Conversely, PCA performed poorly in the admixed family simulation (both trait models) and the real human genotypes with FES traits.

PCA assumes that genetic relatedness is restricted to a low-dimensional subspace, whereas LMM can handle high-dimensional relatedness. Thus, PCA performs well in the admixture simulation, which is explicitly low-dimensional (see Materials and Methods), and our subpopulation tree simulations, which are likely well approximated by a few dimensions despite the large number of subpopulations because there are few long branches. Conversely, PCA performs poorly under family structure because its kinship matrix is high-dimensional (Fig. S7). However, estimating the latent space dimensions of real datasets is challenging because estimated eigenvalues have biased distributions [113]. Kinship matrix rank estimated using the Tracy-Widom test [28] did not fully predict the datasets that PCA performs well on. In contrast, estimated local kinship finds considerable cryptic family relatedness in all real human datasets and better explains why PCA performs poorly there. The trait model also influences the relative performance of PCA, so genotype-only parameters (eigenvalues or local kinship) alone cannot tell the full story. There are related tests for numbers of dimensions that consider the trait which we did not consider, including the Bayesian information criterion for the regression with PCs against the trait [38]. Additionally, PCA and LMM goodness of fit could be compared using the coefficient of determination generalized for LMMs [114]. PCA is at best underpowered relative to LMMs, and at worst miscalibrated regardless of the numbers of PCs included, in real human genotype tests. Among our simulations, such poor performance occurred only in the admixed family. Local kinship estimates reveal considerable family relatedness in the real datasets absent in the corresponding subpopulation tree simulations. Admixture is also absent in our tree simulations, but our simulations and theory show that admixture is well handled by PCA. Hundreds of close relative pairs have been identified in 1000 Genomes [115–118], but their removal does not improve PCA performance sufficiently in our tests, so the larger number of more distantly related pairs are PCA’s most serious obstacle in practice. Distant relatives are expected to be numerous in any large human dataset [77, 119, 120]. Our FES trait tests show that family relatedness is more challenging when rarer variants have larger coefficients. Overall, the high relatedness dimensions induced by family relatedness is the key challenge for PCA association in modern datasets that is readily overcome by LMM.

Our tests also found PCA robust to large numbers of PCs, far beyond the optimal choice, agreeing with previous anecdotal observations [26, 56], in contrast to using too few PCs for which there is a large performance penalty. The exception was the small sample size simulation, where only small numbers of PCs performed well. In contrast, LMM is simpler since there is no need to choose the number of PCs. However, an LMM with a large number of covariates may have conservative p-values, as observed for LMM with large numbers of PCs, which is a weakness of the score test used by the LMM we evaluated that may be overcome with other statistical tests. Simulations or post hoc evaluations remain crucial for ensuring that statistics are calibrated.

There are several variants of the PCA and LMM analyses, most designed for better modeling linkage disequilibrium (LD), that we did not evaluate directly, in which PCs are no longer exactly the top eigenvectors of the kinship matrix (if estimated with different approaches), although this is not a crucial aspect of our arguments. We do not consider the case where samples are projected onto PCs estimated from an external sample [121], which is uncommon in association studies, and whose primary effect is shrinkage, so if all samples are projected then they are all equally affected and larger regression coefficients compensate for the shrinkage, although this will no longer be the case if only a portion of the sample is projected onto the PCs of the rest of the sample. Another approach tests PCs for association against every locus in the genome in order to identify and exclude PCs that capture LD structure (which is localized) instead of ancestry (which should be present across the genome) [121]; a previous proposal removes LD using an autocorrelation model prior to estimating PCs [28]. These improved PCs remain inadequate models of family relatedness, so an LMM will continue to outperform them in that setting. Similarly, the leave-one-chromosome-out (LOCO) approach for estimating kinship matrices for LMMs prevents the test locus and loci in LD with it from being modeled by the random effect as well, which is called “proximal contamination” [55, 62]. While LOCO kinship estimates vary for each chromosome, they continue to model family relatedness, thus maintaining their key advantage over PCA. The LDAK model estimates kinship instead by weighing loci taking LD into account [122]. LD effects must be adjusted for, if present, so in unfiltered data we advise the previous methods be applied. However, in this work, simulated genotypes do not have LD, and the real datasets were filtered to remove LD, so here there is no proximal contamination and LD confounding is minimized if present at all, so these evaluations may be considered the ideal situation where LD effects have been adjusted successfully, and in this setting LMM outperforms PCA. Overall, these alternative PCs or kinship matrices differ from their basic counterparts by either the extent to which LD influences the estimates (which may be a confounder in a small portion of the genome, by definition) or by sampling noise, neither of which are expected to change our key conclusion.

One of the limitations of this work include relatively small sample sizes compared to modern association studies. However, our conclusions are not expected to change with larger sample sizes, as cryptic family relatedness will continue to be abundant in such data, if not increase in abundance, and thus give LMMs an advantage over PCA [77, 119, 120]. One reason PCA has been favored over classic LMMs is because PCA’s runtime scales much better with increasing sample size. However, recent approaches not tested in this work have made LMMs more scalable and applicable to biobank-scale data [59, 67, 72], so one clear next step is carefully evaluating these approaches in simulations with larger sample sizes. A different benefit for including PCs were recently reported for BOLTLMM, which does not result in greater power but rather in reduced runtime, a property that may be specific to its use of scalable algorithms such as conjugate gradient and variational Bayes [77]. Many of these newer LMMs also no longer follow the infinitesimal model of the basic LMM [67, 72], and employ novel approximations, which are features not evaluated in this work and worthy of future study.

Another limitation of this work is ignoring rare variants, a necessity given our smaller sample sizes, where rare variant association is miscalibrated and underpowered. Using simulations mimicking the UK Biobank, recent work has found that rare variants can have a more pronounced structure than common variants, and that modeling this rare variant structure (with either PCA and LMM) may better model environment confounding, reduce inflation in association studies, and ameliorate stratification in polygenic risk scores [123]. Better modeling rare variants and their structure is a key next step in association studies.

The largest limitation of our work is that we only considered quantitative traits. Previous evaluations involving case-control traits tended to report PCA-LMM ties or mixed results, an observation potentially confounded by the use of low-dimensional simulations without family relatedness (Table 1). An additional concern is case-control ascertainment bias and imbalance, which appears to affect LMMs more severely, although recent work appears to solve this problem [55, 59]. Future evaluations should aim to include our simulations and real datasets, to ensure that previous results were not biased in favor of PCA by not simulating family structure or larger coefficients for rare variants that are expected for diseases by various selection models.

Overall, our results lead us to recommend LMM over PCA for association studies in general. Although PCA offer flexibility and speed compared to LMM, additional work is required to ensure that PCA is adequate, including removal of close relatives (lowering sample size and wasting resources) followed by simulations or other evaluations of statistics, and even then PCA may perform poorly in terms of both type I error control and power. The large numbers of distant relatives expected of any real dataset all but ensures that PCA will perform poorly compared to LMM [77, 119, 120]. Our findings also suggest that related applications such as polygenic models may enjoy gains in power and accuracy by employing an LMM instead of PCA to model relatedness [42, 110]. PCA remains indispensable across population genetics, from visualizing population structure and performing quality control to its deep connection to admixture models, but the time has come to limit its use in association testing in favor of LMM or other, richer models capable of modeling all forms of relatedness.

## Abbreviations

PCA: principal component analysis;
PCs: principal components;
LMM: linear mixed-effects model;
FES: fixed effect sizes (trait model);
RC: random coefficients (trait model);
MAF: minor allele frequency;
WGS: whole genome sequencing;
LD: linkage disequilibrium.

## 5 Appendices

### 5.1 Appendix A: Fitting ancestral allele frequency distribution to real data

We calculated 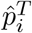 distributions of each real dataset. However, population structure increases the variance of these sample 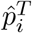 relative to the true 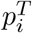 [52]. We present a new algorithm for constructing a new distribution based on the input data but with the lower variance of the true ancestral distribution. Suppose the 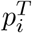 distribution over loci *i* satisfies 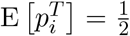 and Var 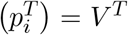. The sample allele frequency 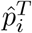, conditioned on 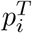, satisfies

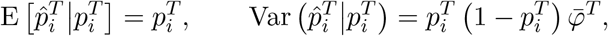

where 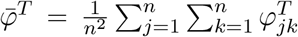 is the mean kinship over all individual [52]. The unconditional moments of 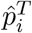 follow from the laws of total expectation and variance: 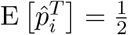 and

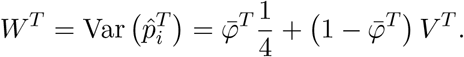

Since 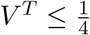 and 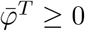, then *W*^*T*^ ≥ *V* ^*T*^. Thus, the goal is to construct a new distribution with the original, lower variance of

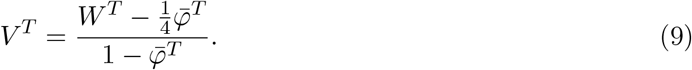

We use the unbiased estimator 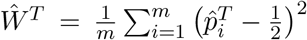, while 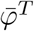 is calculated from the tree parameters: the subpopulation coancestry matrix (Eq. (7)), expanded from subpopulations to individuals, the diagonal converted to kinship (reversing Eq. (8)), and the matrix averaged. However, since our model ignores the MAF filters imposed in our simulations, 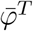 was adjusted. For Human Origins the true model 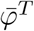 of 0.143 was used. For 1000 Genomes and HGDP the true 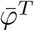 are 0.126 and 0.124, respectively, but 0.4 for both produced a better fit.

Lastly, we construct new allele frequencies,

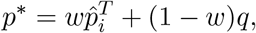

by a weighted average of 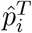 and *q* ∈ (0, 1) drawn independently from a different distribution. 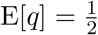 is required to have 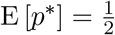. The resulting variance is

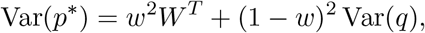

which we equate to the desired *V* ^*T*^ (Eq. (9)) and solve for *w*. For simplicity, we also set Var(*q*) = *V* ^*T*^, which is achieved with:

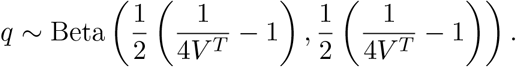

Although *w* = 0 yields Var(*p*^*^) = *V* ^*T*^, we use the second root of the quadratic equation to use 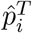:

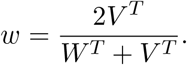

#### 5.2 Appendix B: comparisons between SRMSD_*p*_, AUC_PR_, and evaluation measures from the literature

##### 5.2.1 The inflation factor *λ*

Test statistic inflation has been used to measure model calibration [1, 44]. The inflation factor *λ* is defined as the median *χ*^2^ association statistic divided by theoretical median under the null hypothesis [23]. To compare p-values from non-*χ*^2^ tests (such as t-statistics), *λ* can be calculated from p-values using

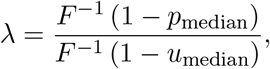

where *p*_median_ is the median observed p-value (including causal loci), 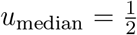 is its null expectation, and *F* is the *χ*^2^ cumulative density function (*F*^−1^ is the quantile function).

To compare *λ* and SRMSD_*p*_ directly, for simplicity assume that all p-values are null. In this case, calibrated p-values give *λ* = 1 and SRMSD_*p*_ = 0. However, non-uniform p-values with the expected median, such as from genomic control [23], result in *λ* = 1, but SRMSD_*p*_*≠* 0 except for uniform p-values, a key flaw of *λ* that SRMSD_*p*_ overcomes. Inflated statistics (anti-conservative p-values) give *λ >* 1 and SRMSD_*p*_ > 0. Deflated statistics (conservative p-values) give *λ* < 1 and SRMSD_*p*_ < 0. Thus, *λ≠* 1 always implies SRMSD_*p*_*≠* 0 (where *λ* − 1 and SRMSD_*p*_ have the same sign), but not the other way around. Overall, *λ* depends only on the median p-value, while SRMSD_*p*_ uses the complete distribution. However, SRMSD_*p*_ requires knowing which loci are null, so unlike *λ* it is only applicable to simulated traits.

##### 5.2.2 Empirical comparison of SRMSD_*p*_ and *λ*

There is a near one-to-one correspondence between *λ* and SRMSD_*p*_ in our data (Fig. S1). PCA tended to be inflated (*λ >* 1 and SRMSD_*p*_ > 0) whereas LMM tended to be deflated (*λ* < 1 and SRMSD_*p*_ < 0), otherwise the data for both models fall on the same contiguous curve. We fit a sigmoidal function to this data,

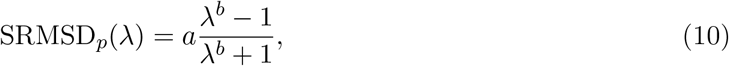

which for *a, b >* 0 satisfies SRMSD_*p*_(*λ* = 1) = 0 and reflects log(*λ*) about zero (*λ* = 1): SRMSD_*p*_(log(*λ*) = −*x*) = −SRMSD_*p*_(log(*λ*) = *x*).

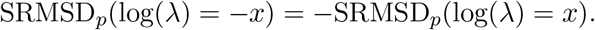

We fit this model to *λ >* 1 only since it was less noisy and of greater interest, and obtained the curve shown in Fig. S1 with *a* = 0.564 and *b* = 0.619. The value *λ* = 1.05, a common threshold for benign inflation [44], corresponds to SRMSD_*p*_ = 0.0085 according to Eq. (10). Conversely, SRMSD_*p*_ = 0.01, serving as a simpler rule of thumb, corresponds to *λ* = 1.06.

##### 5.2.3 Type I error rate

The type I error rate is the proportion of null p-values with *p* ≤ *t*. Calibrated p-values have type I error rate near *t*, which may be evaluated with a binomial test. This measure may give different results for different *t*, for example be significantly miscalibrated only for large *t* (due to lack of power for smaller *t*), and it requires large simulations to estimate well as it depends on the tail of the distribution. In contrast, SRMSD_*p*_ uses the entire distribution so it is easier to estimate, SRMSD_*p*_ = 0 guarantees calibrated type I error rates at all *t*, while large |SRMSD_*p*_| indicates incorrect type I errors for a range of *t*. Empirically, we find the expected agreement and monotonic relationship between SRMSD_*p*_ and type I error rate (Fig. S2).

##### 5.2.4 Statistical power and comparison to AUC_PR_

Power is the probability that a test is declared significant when the alternative hypothesis *H*_1_ holds. At a p-value threshold *t*, power equals

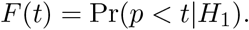

*F* (*t*) is a cumulative function, so it is monotonically increasing and has an inverse. Like type I error control, power may rank models differently depending on *t*, and it is also harder to estimate than AUC_PR_ because power depends on the tail of the distribution.

Power is not meaningful when p-values are not calibrated. To establish a clear connection to AUC_PR_, assume calibrated (uniform) null p-values: Pr(*p < t*|*H*_0_) = *t*. TPs, FPs, and FNs at *t* are

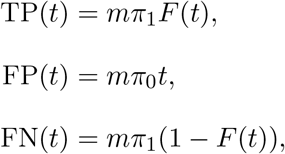

where *π*_0_ = Pr(*H*_0_) is the proportion of null cases and *π*_1_ = 1 − *π*_0_ of alternative cases. Therefore,

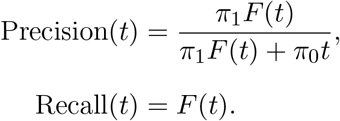

Noting that *t* = *F*^−1^(Recall), precision can be written as a function of recall, the power function, and constants:

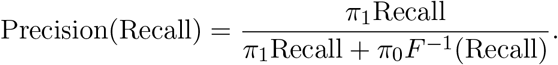

This last form leads most clearly to 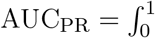 Precision(Recall)*d*Recall.

Lastly, consider a simple yet common case in which model *A* is uniformly more powerful than model *B*: *F*_*A*_(*t*) *> F*_*B*_(*t*) for every *t*. Therefore 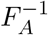 (Recall) < 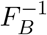 (Recall) for every recall value. This ensures that the precision of *A* is greater than that of *B* at every recall value, so AUC_PR_ is greater for *A* than *B*. Thus, AUC_PR_ ranks calibrated models according to power.

Empirically, we find the predicted positive correlation between AUC_PR_ and calibrated power (Fig. S3). The correlation is clear when considered separately per dataset, but the slope varies per dataset, which is expected because the proportion of alternative cases *π*_1_ varies per dataset.

### Competing interests

The authors declare no competing interests.

## Acknowledgments

Thanks to Tiffany Tu, Ratchanon Pornmongkolsuk, and Zhuoran Hou for feedback on this article. This work was funded in part by the Duke University School of Medicine Whitehead Scholars Program, a gift from the Whitehead Charitable Foundation. The 1000 Genomes data were generated at the New York Genome Center with funds provided by NHGRI Grant 3UM1HG008901-03S1.

## Web resources

plink2, https://www.cog-genomics.org/plink/2.0/

GCTA, https://yanglab.westlake.edu.cn/software/gcta/

Eigensoft, https://github.com/DReichLab/EIG

bnpsd, https://cran.r-project.org/package=bnpsd

simfam, https://cran.r-project.org/package=simfam

simtrait, https://cran.r-project.org/package=simtrait

genio, https://cran.r-project.org/package=genio

popkin, https://cran.r-project.org/package=popkin

ape, https://cran.r-project.org/package=ape

nnls, https://cran.r-project.org/package=nnls

PRROC, https://cran.r-project.org/package=PRROC

BEDMatrix, https://cran.r-project.org/package=BEDMatrix

## Data and code availability

The data and code generated during this study are available on GitHub at https://github.com/OchoaLab/pca-assoc-paper. The public subset of Human Origins is available on the Reich Lab website at https://reich.hms.harvard.edu/datasets; non-public samples have to be requested from David Reich. The WGS version of HGDP was downloaded from the Wellcome Sanger Institute FTP site at ftp://ngs.sanger.ac.uk/production/hgdp/hgdp_wgs.20190516/. The highcoverage version of the 1000 Genomes Project was downloaded from ftp://ftp.1000genomes.ebi.ac.uk/vol1/ftp/data_collections/1000G_2504_high_coverage/working/20190425_NYGC_GATK/.

## Supplemental figures

**Figure S1:**
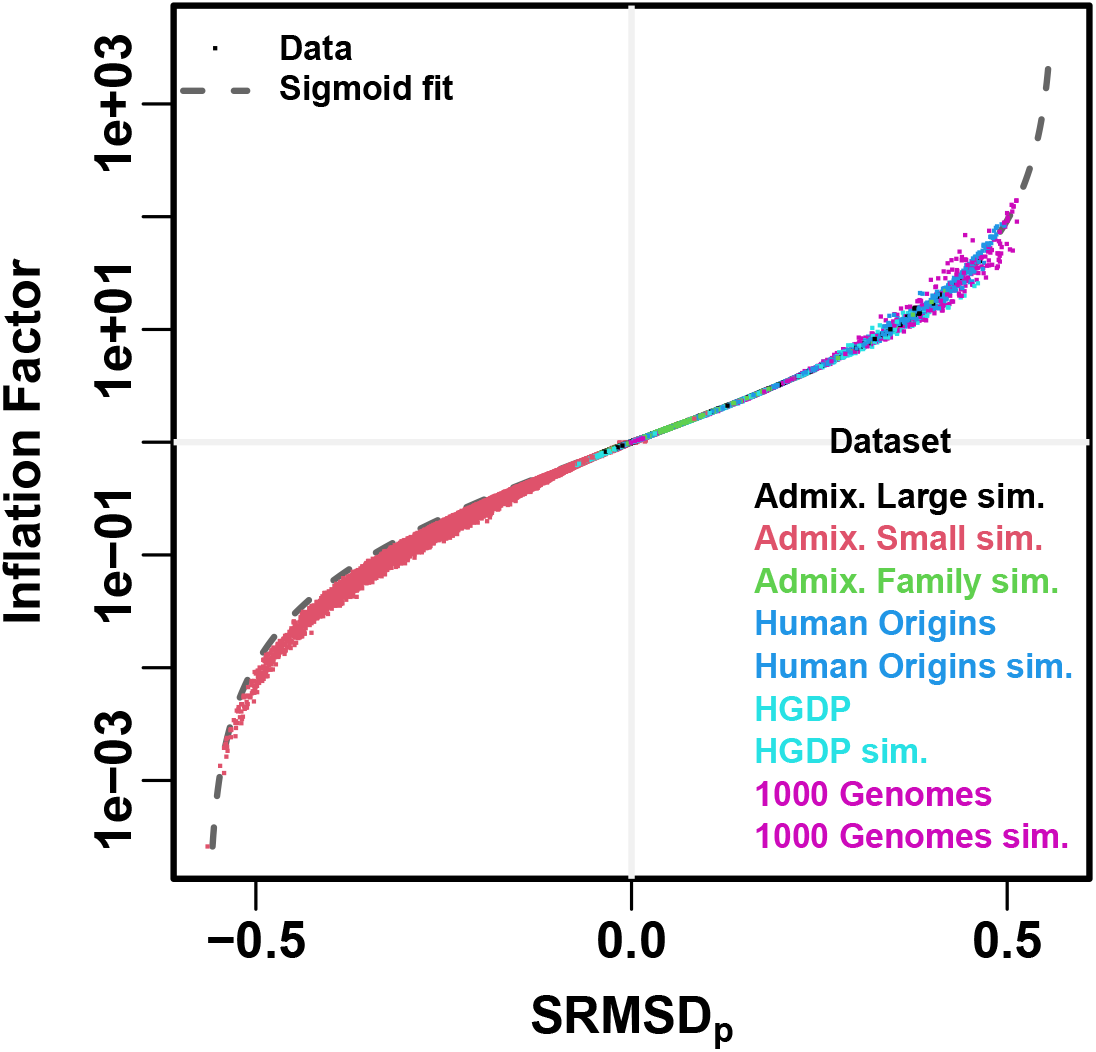
Comparison between SRMSD_*p*_ and inflation factor. Each point is a pair of statistics for one replicate, one association model (PCA or LMM with some number of PCs *r*), one trait model (FES vs RC, all heritability/environments tested), and one dataset (color coded by dataset). Note log y-axis. The sigmoidal curve in Eq. (10) is fit to the data.

**Figure S2:**
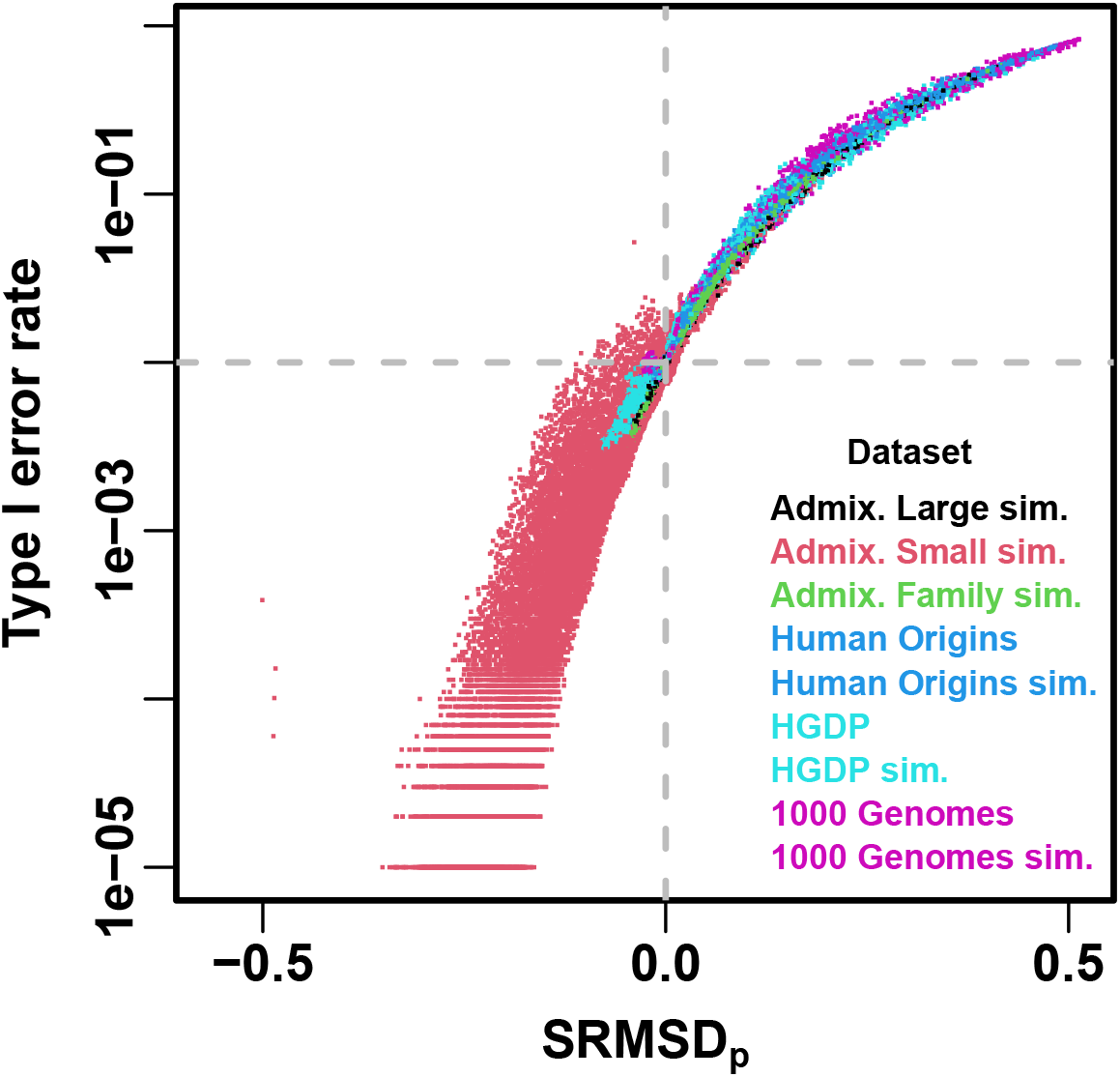
Comparison between SRMSD_*p*_ and type I error rate. Type I error rate calculated at a p-value threshold of 1e-2 (horizontal dashed gray line). Thus, a calibrated model has a type I error rate of 1e-2 and SRMSD_*p*_ = 0 (where the dashed lines meet). As expected, increased type I error rates correspond to SRMSD_*p*_ > 0, while reduced type I error rates correspond to SRMSD_*p*_ < 0. Each point is a pair of statistics for one replicate, one association model (PCA or LMM with some number of PCs *r*), one trait model (FES vs RC, all heritability/environments tested), and one dataset (color coded by dataset). Note log y-axis.

**Figure S3:**
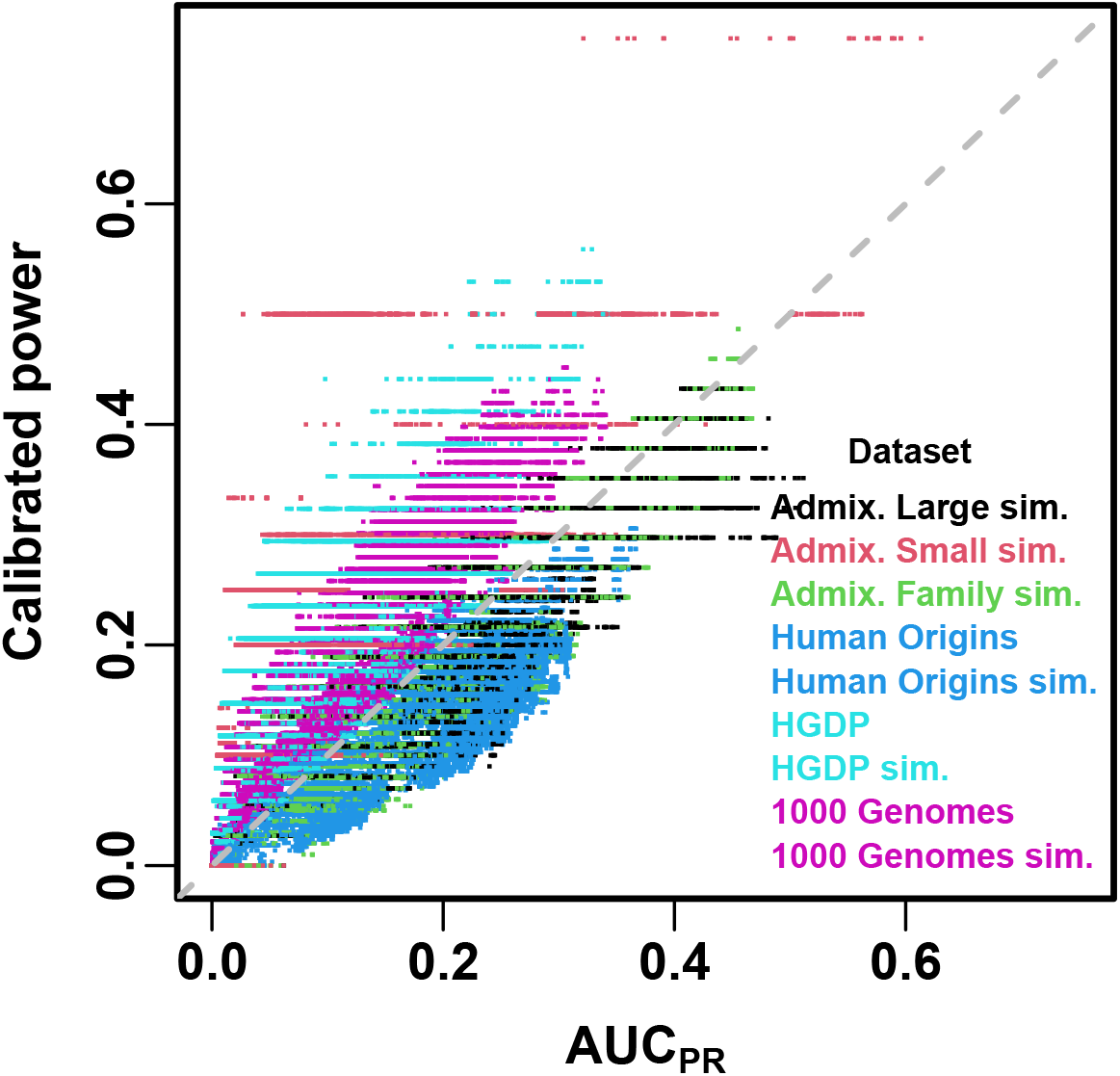
Comparison between AUC_PR_ and calibrated power. Calibrated power is power calculated at an empirical type I error threshold of 1e-4. Each point is a pair of statistics for one replicate, one association model (PCA or LMM with some number of PCs *r*), one trait model (FES vs RC, all heritability/environments tested), and one dataset (color coded by dataset). Gray dashed line is *y* = *x* line.

**Figure S4:**
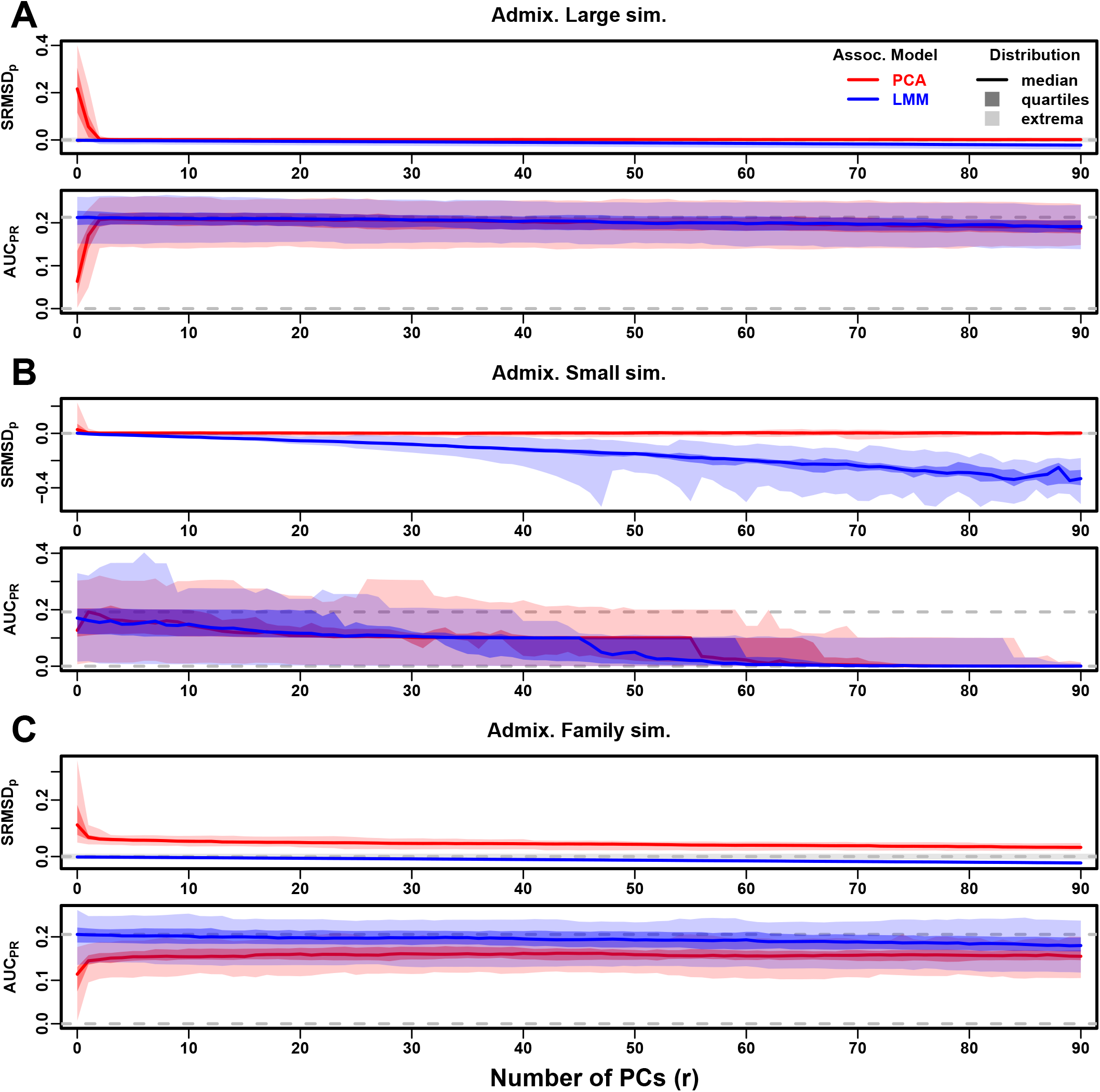
Evaluations in admixture simulations with RC traits. Traits simulated from RC model, otherwise the same as Fig. 3.

**Figure S5:**
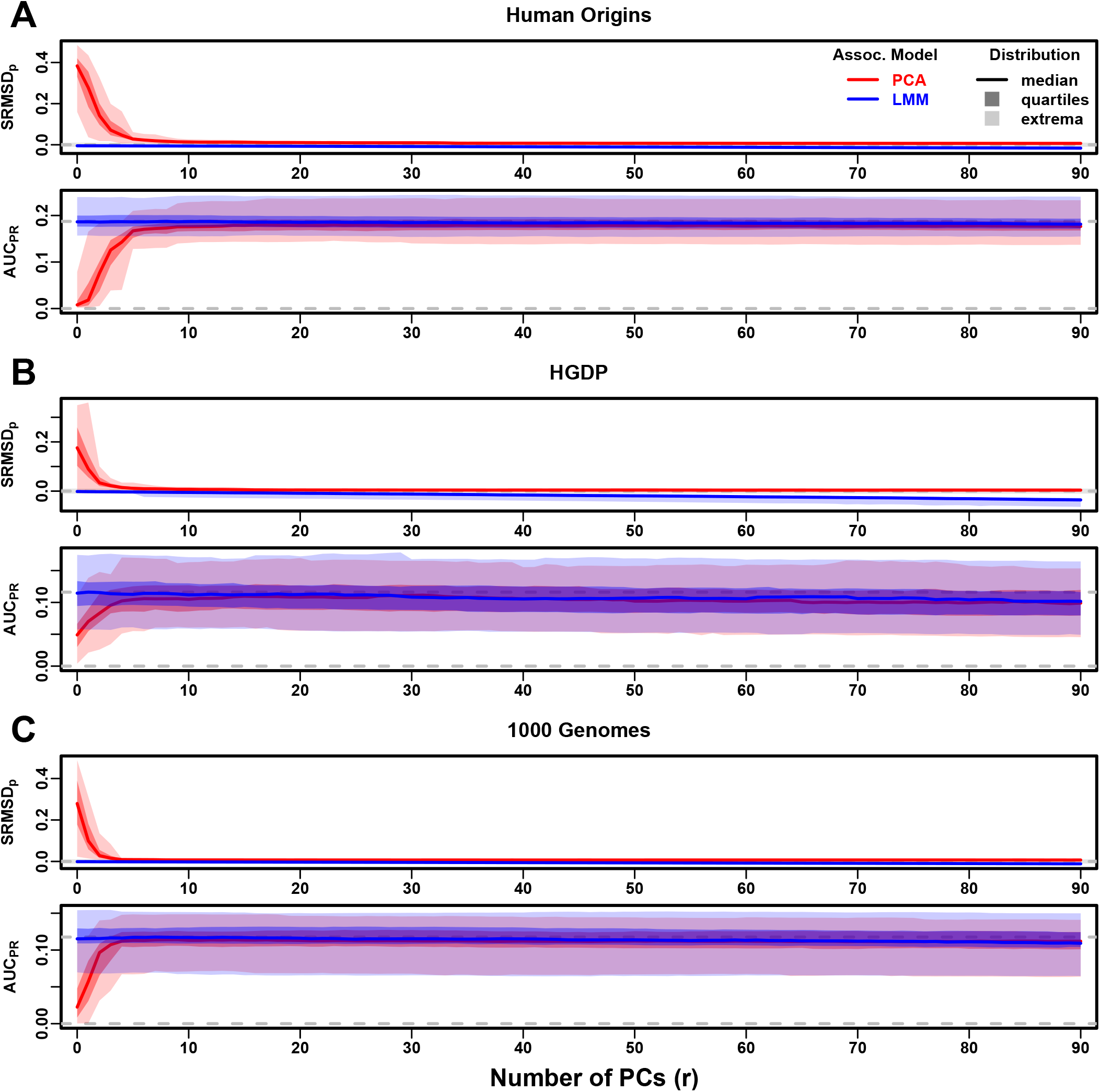
Evaluations in real human genotype datasets with RC traits. Traits simulated from RC model, otherwise the same as Fig. 4.

**Figure S6:**
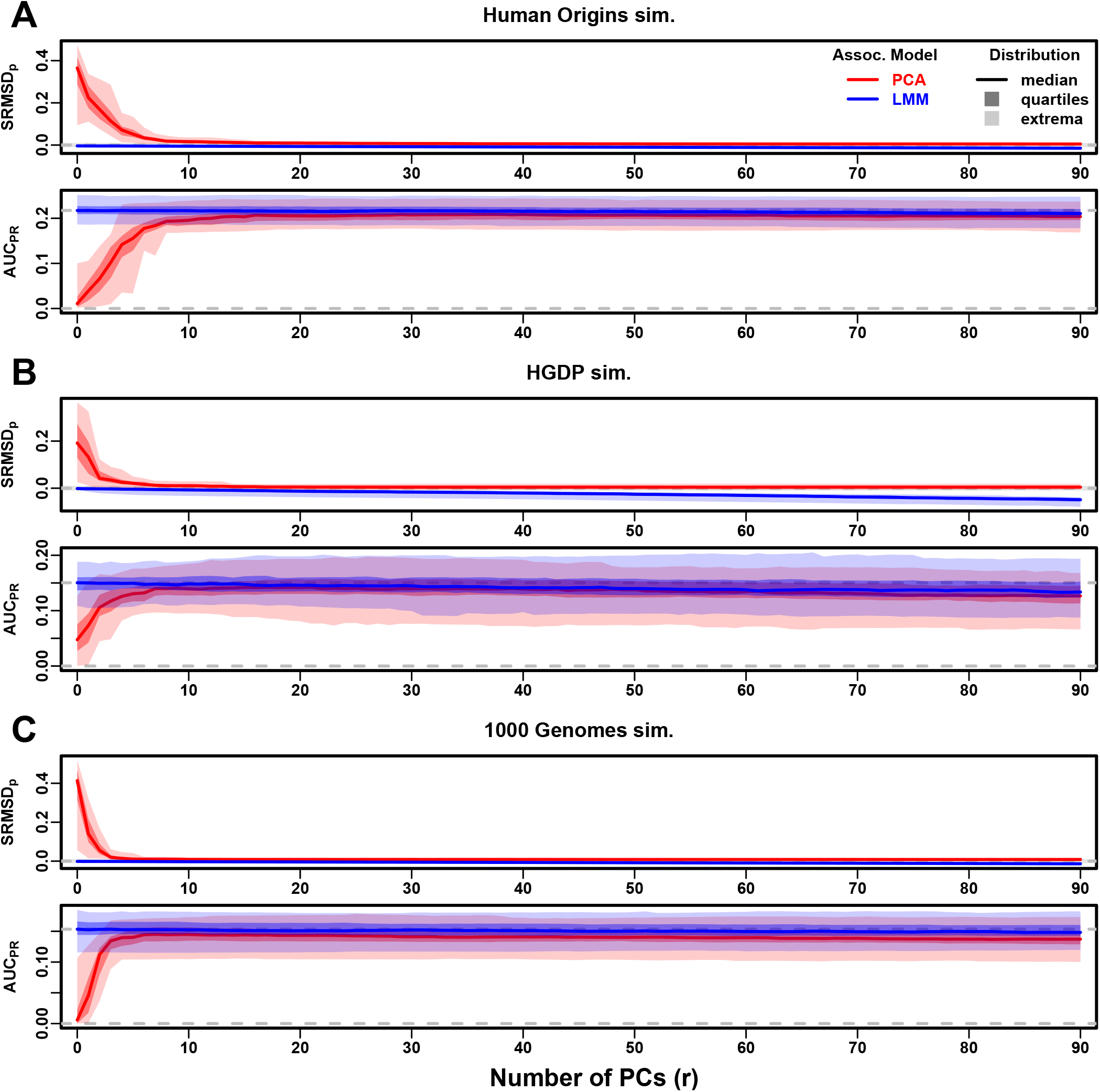
Evaluations in subpopulation tree simulations fit to human data with RC traits. Traits simulated from RC model, otherwise the same as Fig. 5.

**Figure S7:**
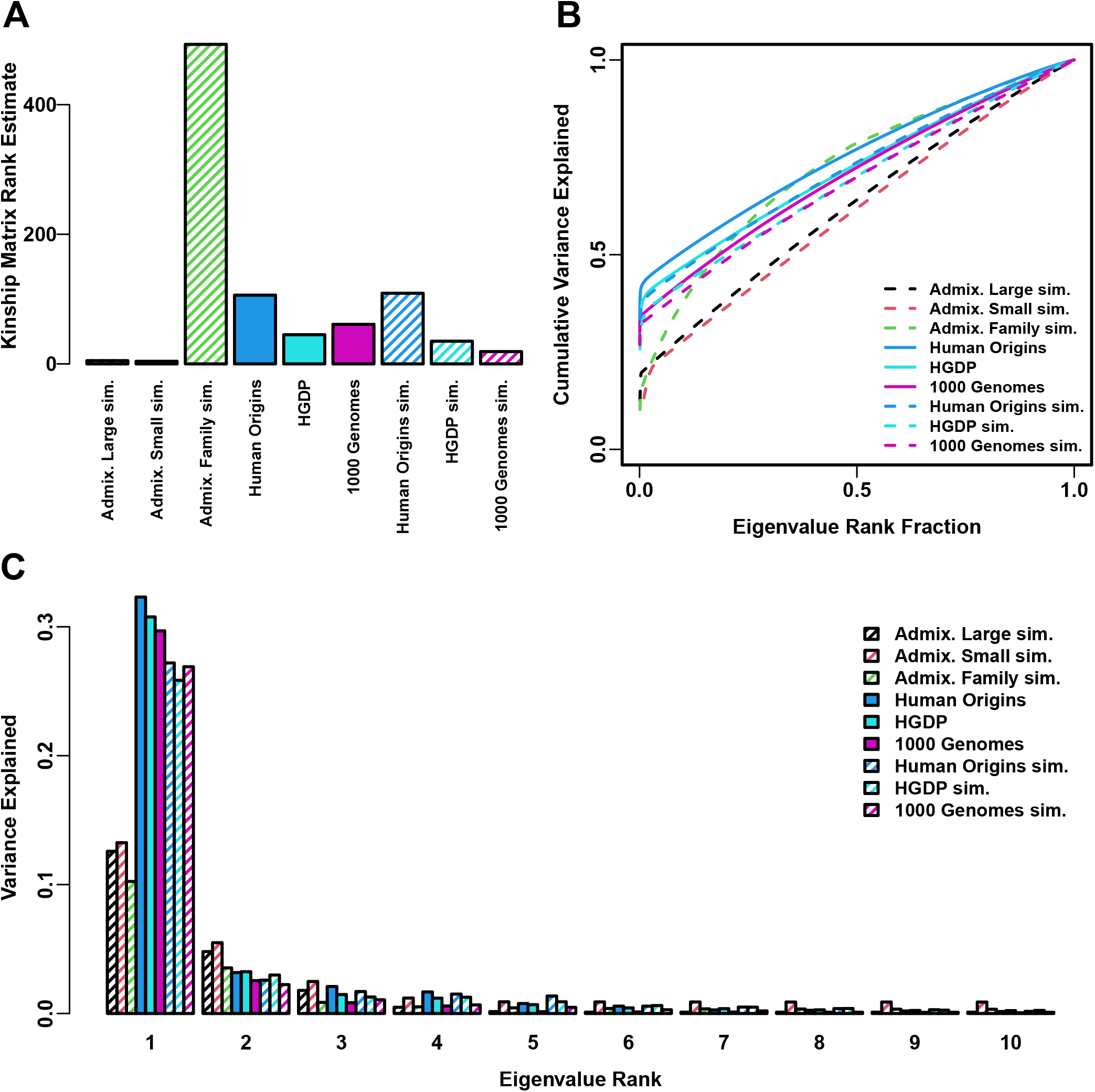
Estimated relatedness dimensions of datasets. **A**. Kinship matrix rank estimated with the Tracy-Widom test with *p* < 0.01. **B**. Cumulative variance explained versus eigenvalue rank fraction. **C**. Variance explained by first 10 eigenvalues.

**Figure S8:**
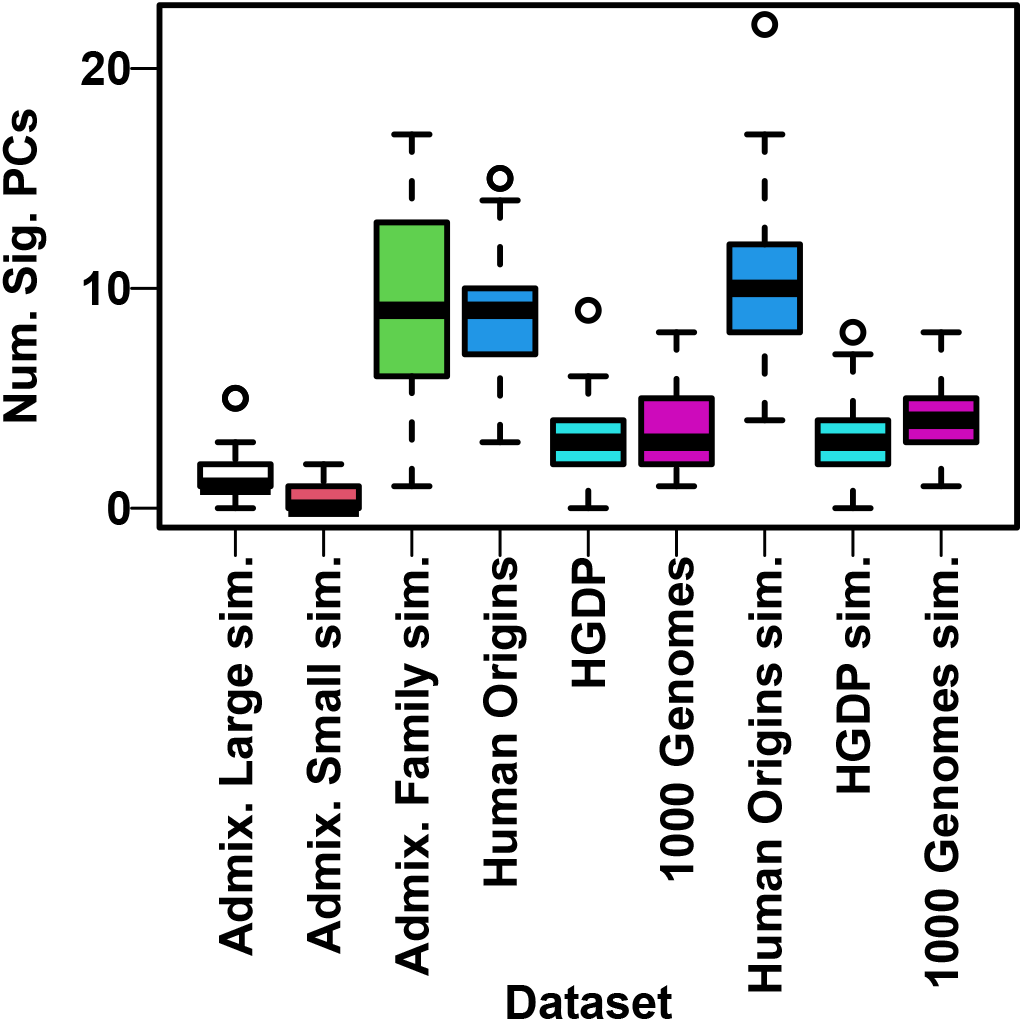
Number of PCs significantly associated with traits. PCs are tested using an ordinary linear regression sequentially, with the *k*th PC tested conditionally on the previous *k* 1− PCs and the intercept. Q-values are estimated from the 90 p-values (one for each PC in a given dataset and replicate) using the R package `qvalue` assuming *π*_0_ = 1 (necessary since the default *π*_0_ estimates were unreliable for such small numbers of p-values and occasionally produced errors), and an FDR threshold of 0.05 is used to determine the number of significant PCs. Distribution per dataset is over its 50 replicates. Shown are results for FES traits with *h*^2^ = 0.8 (the results for RC were very similar, not shown).

**Figure S9:**
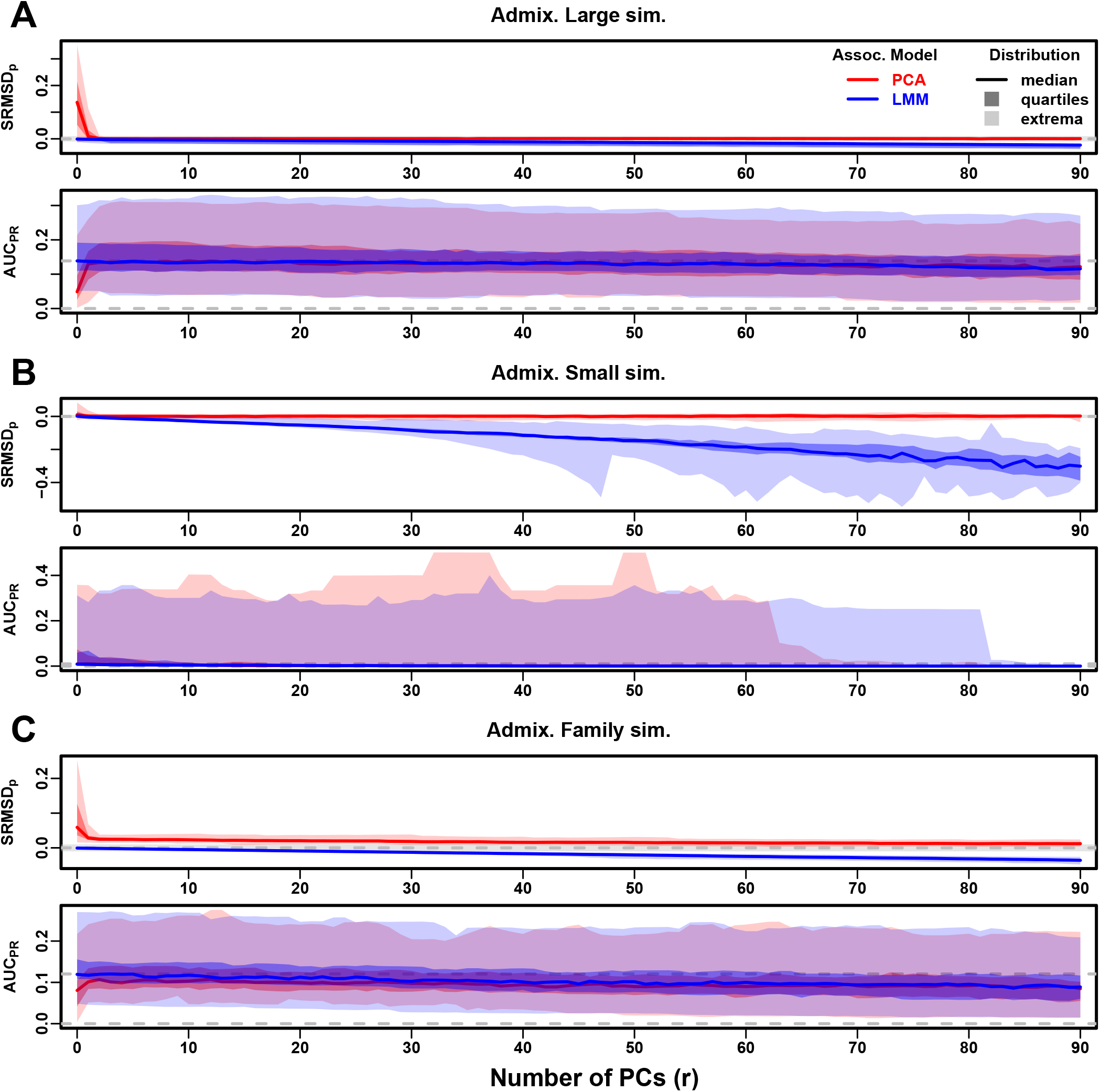
Evaluations in admixture simulations with FES traits, low heritability. Traits simulated using *h*^2^ = 0.3, otherwise the same as Fig. 3.

**Figure S10:**
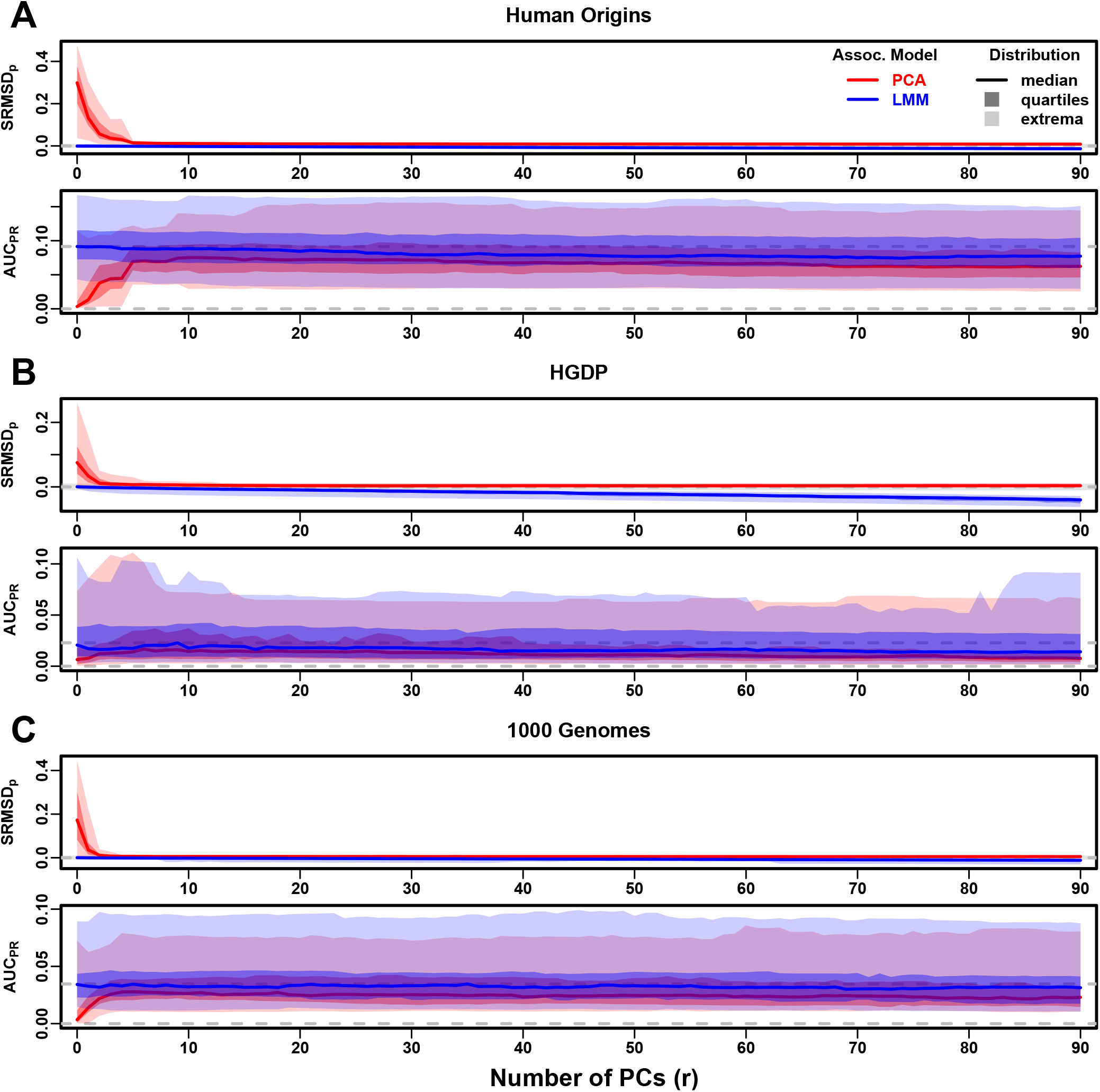
Evaluations in real human genotype datasets with FES traits, low heritability. Traits simulated using *h*^2^ = 0.3, otherwise the same as Fig. 4.

**Figure S11:**
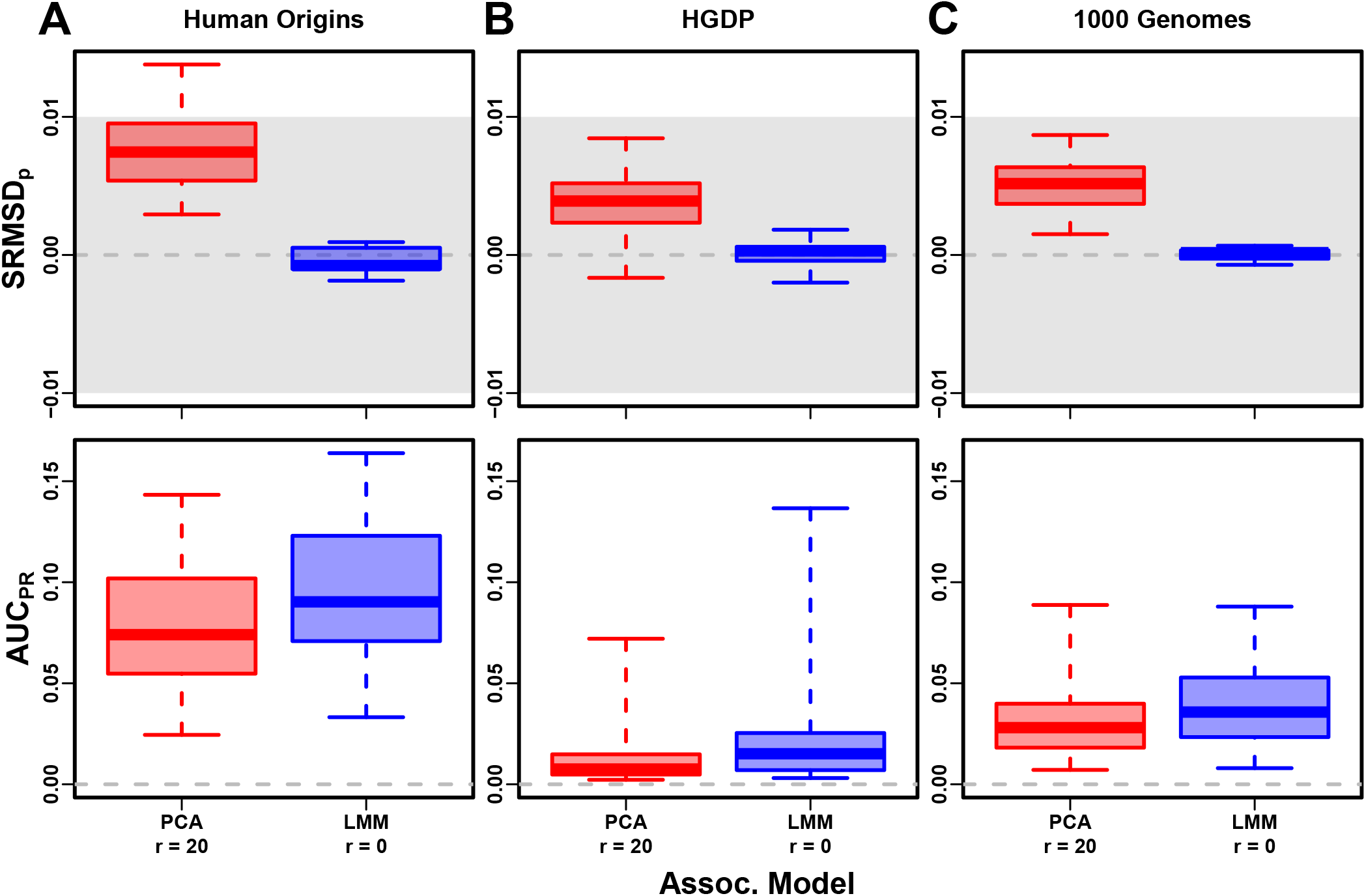
Evaluation in real datasets excluding 4th degree relatives, FES traits, low heritability. Traits simulated using *h*^2^ = 0.3, otherwise the same as Fig. 7.

**Figure S12:**
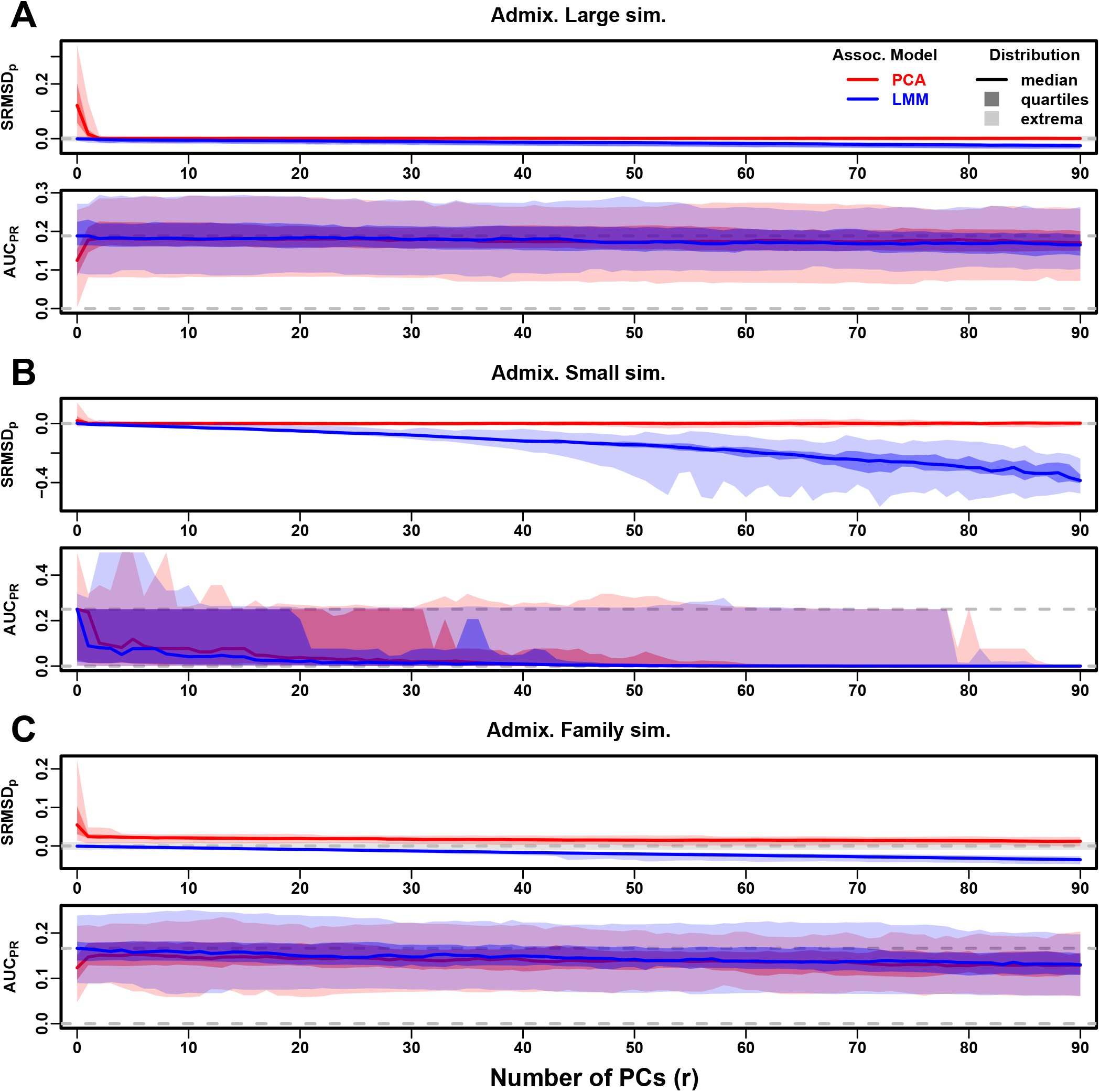
Evaluations in admixture simulations with RC traits, low heritability. Traits simulated using *h*^2^ = 0.3, otherwise the same as Fig. S4.

**Figure S13:**
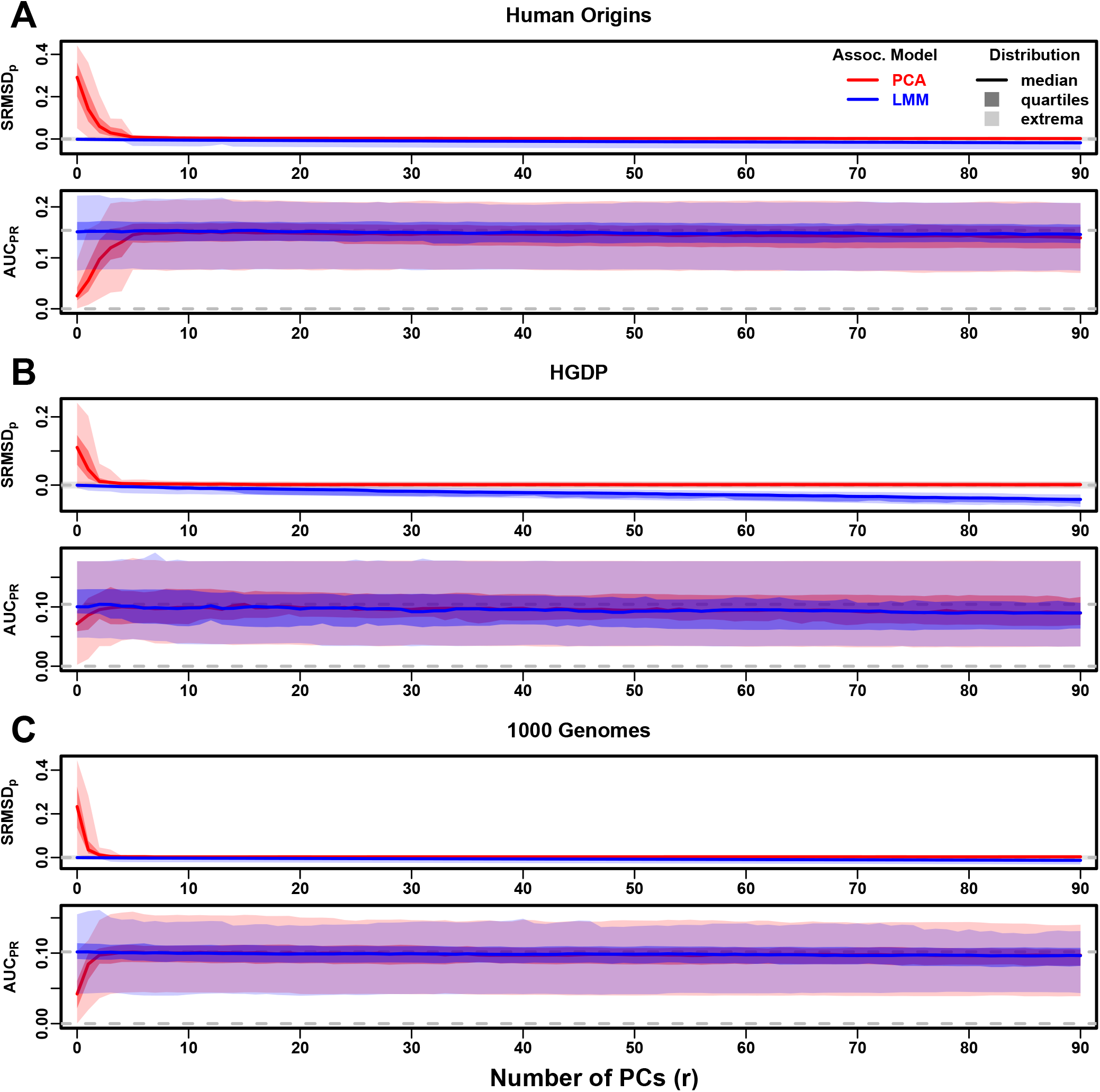
Evaluations in real human genotype datasets with RC traits, low heritability. Traits simulated using *h*^2^ = 0.3, otherwise the same as Fig. S5.

**Figure S14:**
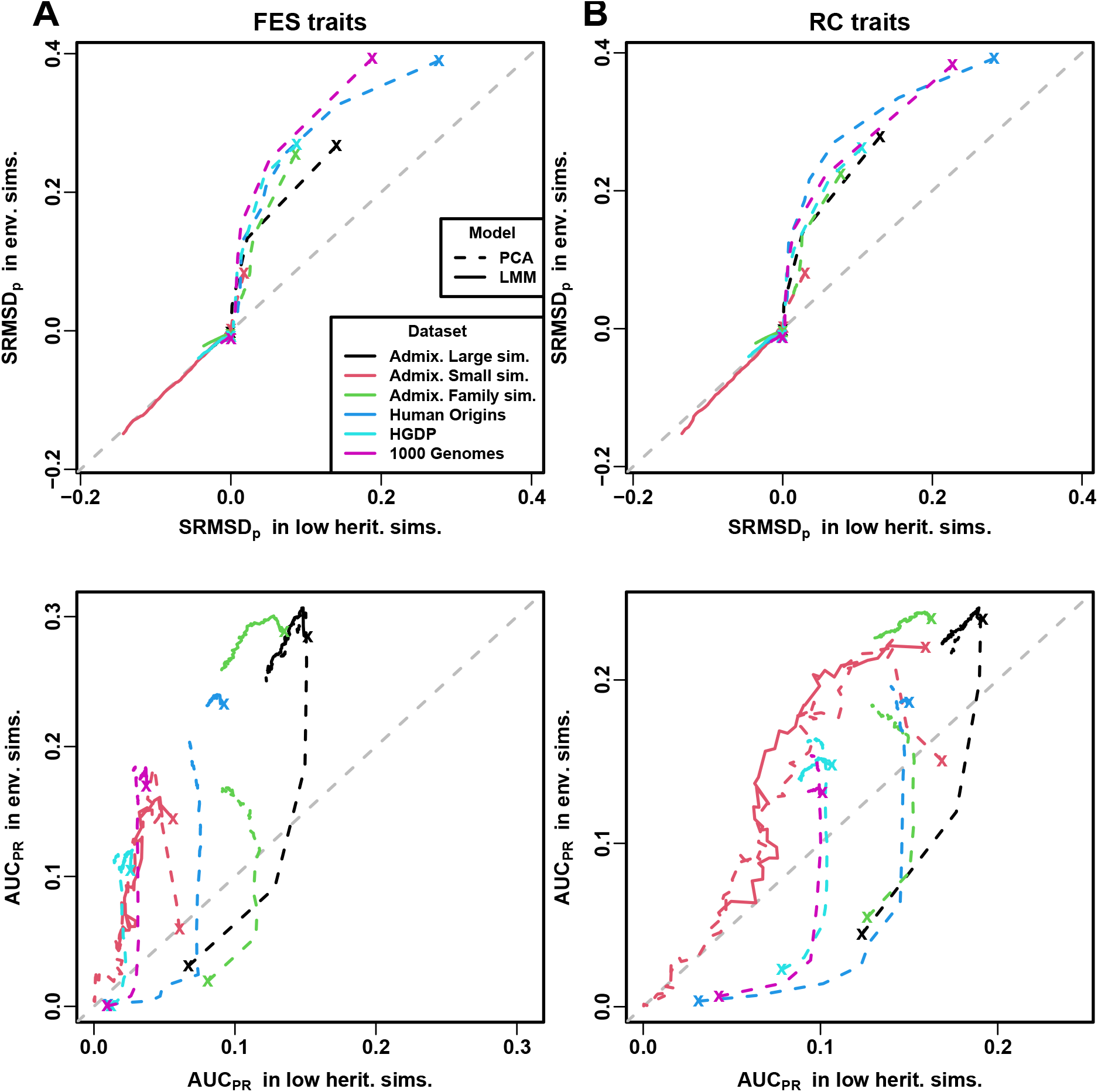
Comparison of performance in low heritability vs environment simulations. Each curve traces as the number of PCs *r* is increased from *r* = 0 (marked with an “x”) until *r* = 90 (unmarked end), on one axis is the mean value over replicates of either SRMSD_*p*_ or AUC_PR_, for low heritability simulations on the x-axis and environment simulations on the y-axis. Each curve corresponds to one dataset (color) and association model (solid or dashed line type). Columns: **A**. FES and **B**. RC traits show similar results. First row shows that for PCA curves (dashed), SRMSD_*p*_ is higher (worse) in environment simulations for low *r*, but becomes equal in both simulations once *r* is sufficiently large; for LMM curves (solid), SRMSD_*p*_ is equal in both simulations for all *r*, all datasets. Second row shows that for PCA, AUC_PR_ is higher (better) in low heritability simulations for low *r*, but becomes higher in environment simulations once *r* is sufficiently large; for LMM, performance is better in environment simulations for all *r*, all datasets.

**Figure S15:**
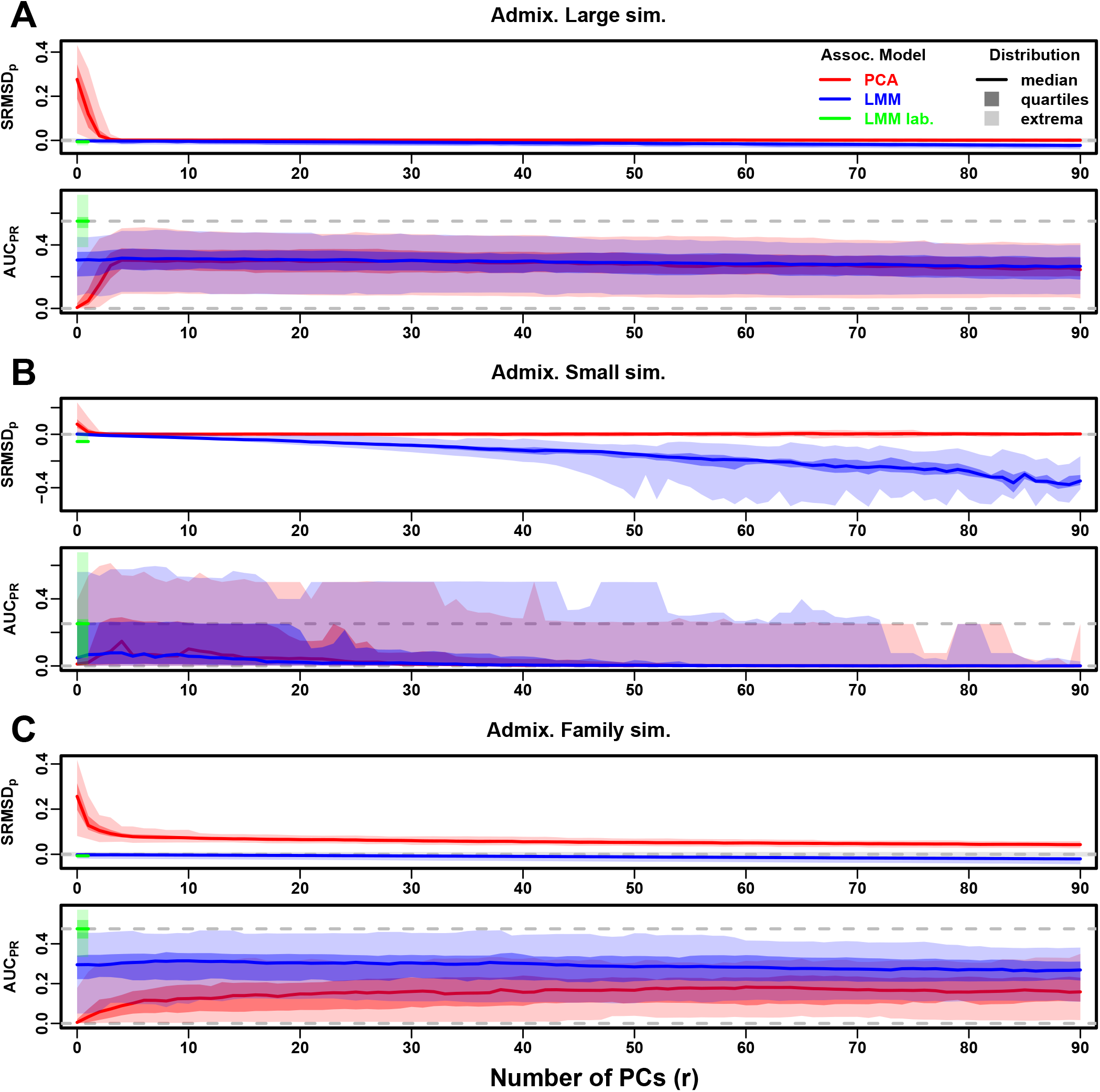
Evaluations in admixture simulations with FES traits, environment. Traits simulated with environment effects, otherwise the same as Fig. S9. “LMM lab.” was only tested with *r* = 0.

**Figure S16:**
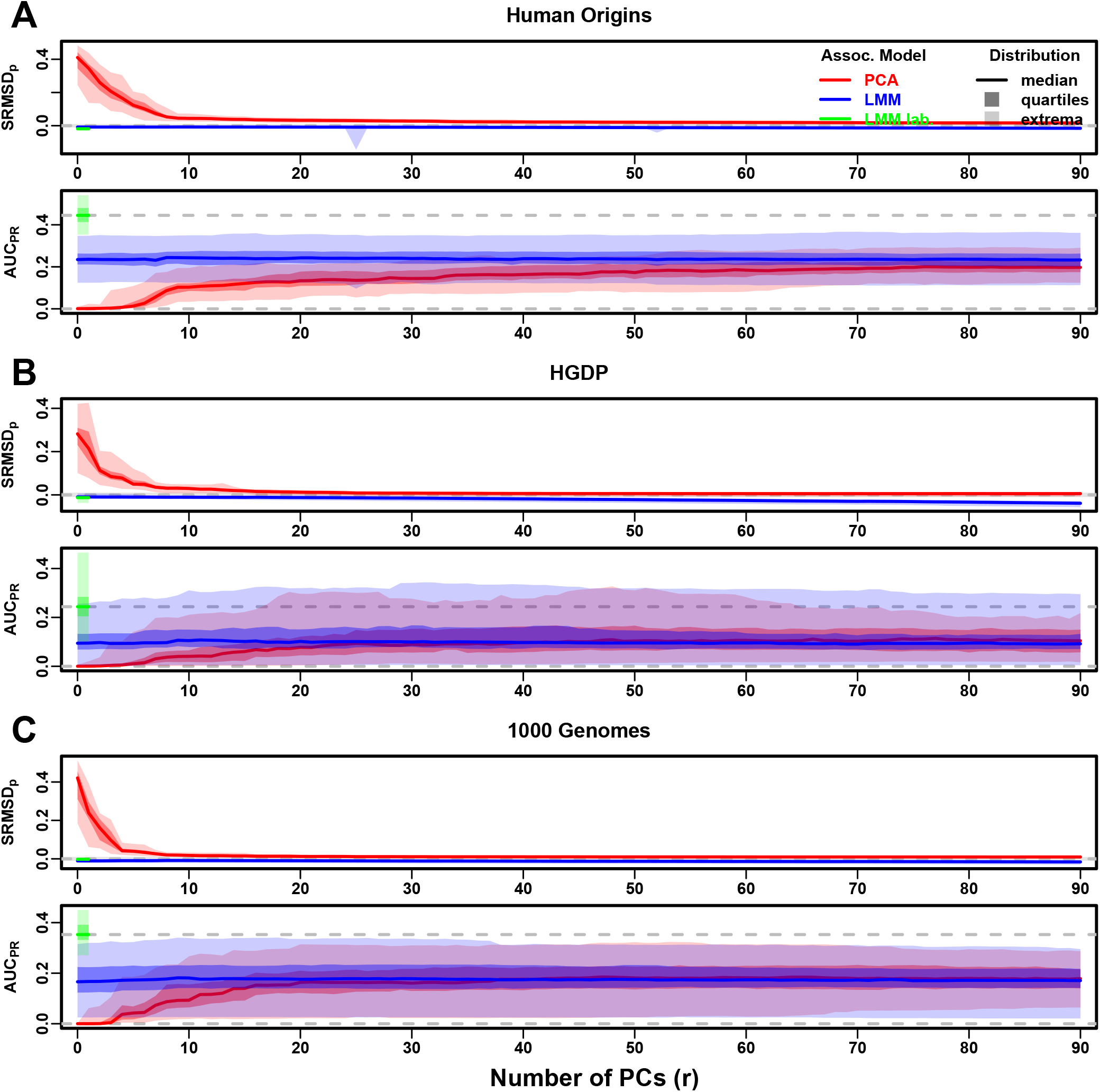
Evaluations in real human genotype datasets with FES traits, environment. Traits simulated with environment effects, otherwise the same as Fig. S10. “LMM lab.” was only tested with *r* = 0.

**Figure S17:**
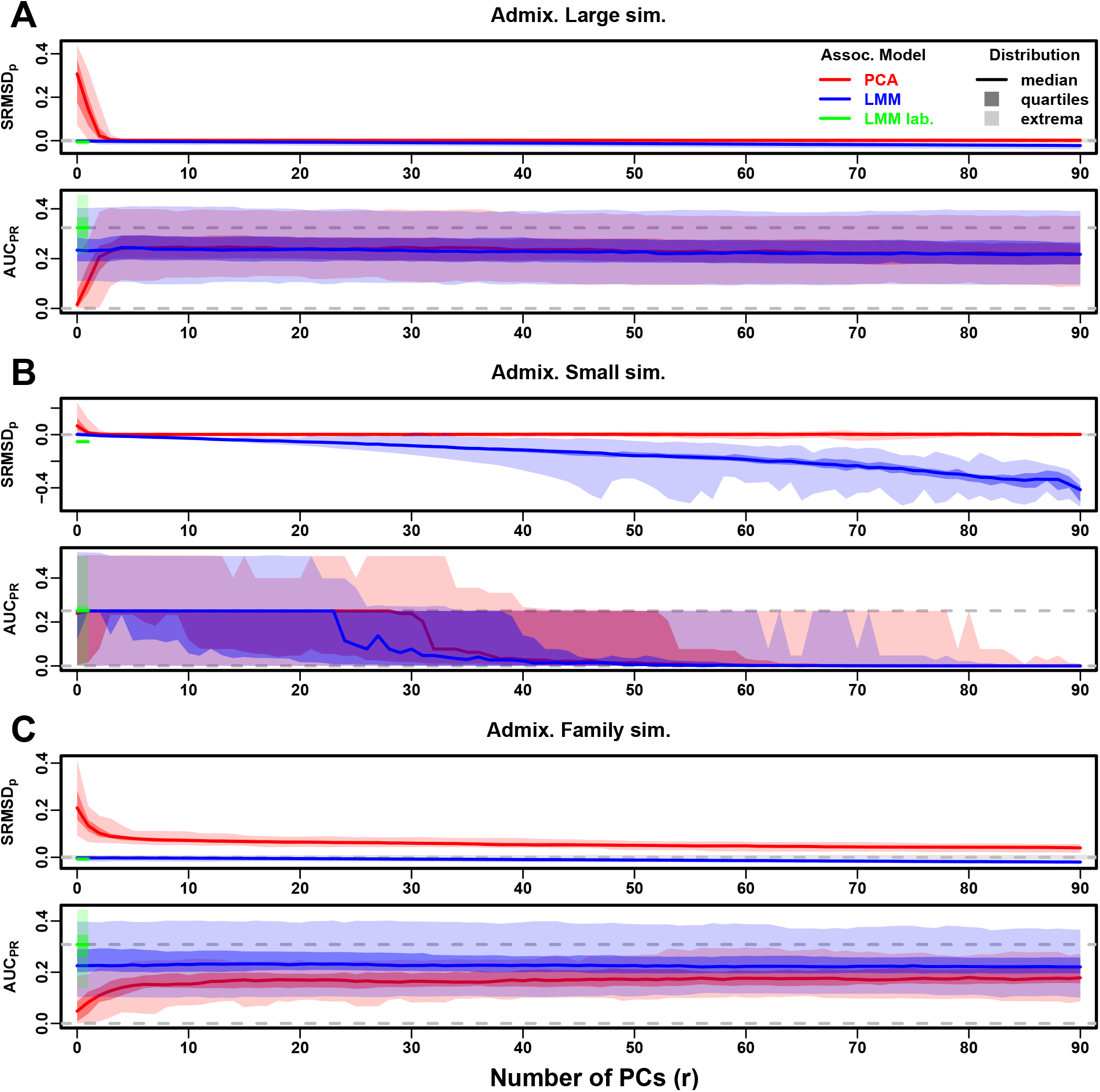
Evaluations in admixture simulations with RC traits, environment. Traits simulated with environment effects, otherwise the same as Fig. S12. “LMM lab.” was only tested with *r* = 0.

**Figure S18:**
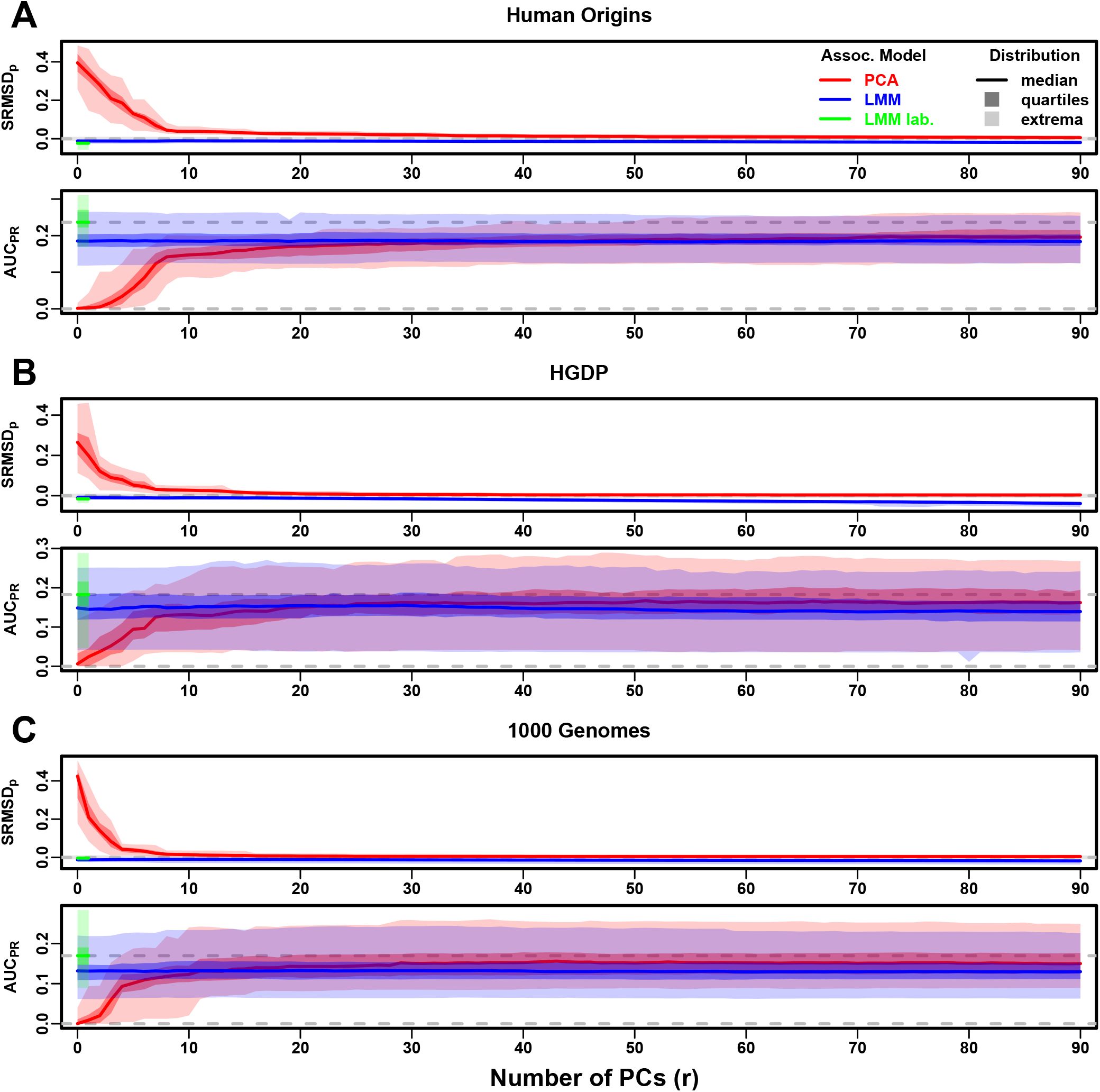
Evaluations in real human genotype datasets with RC traits, environment. Traits simulated with environment effects, otherwise the same as Fig. S13. “LMM lab.” was only tested with *r* = 0.

## Supplemental tables

**Table S1:** Dataset sizes after 4th degree relative filter.

| Dataset | Loci ($m$) | Ind. ($n$) | Ind. removed (%) |
| --- | --- | --- | --- |
| Human Origins | 189,722 | 2636 | 9.8 |
| HGDP | 758,009 | 847 | 8.8 |
| 1000 Genomes | 1,097,415 | 2390 | 4.6 |

**Table S2:**
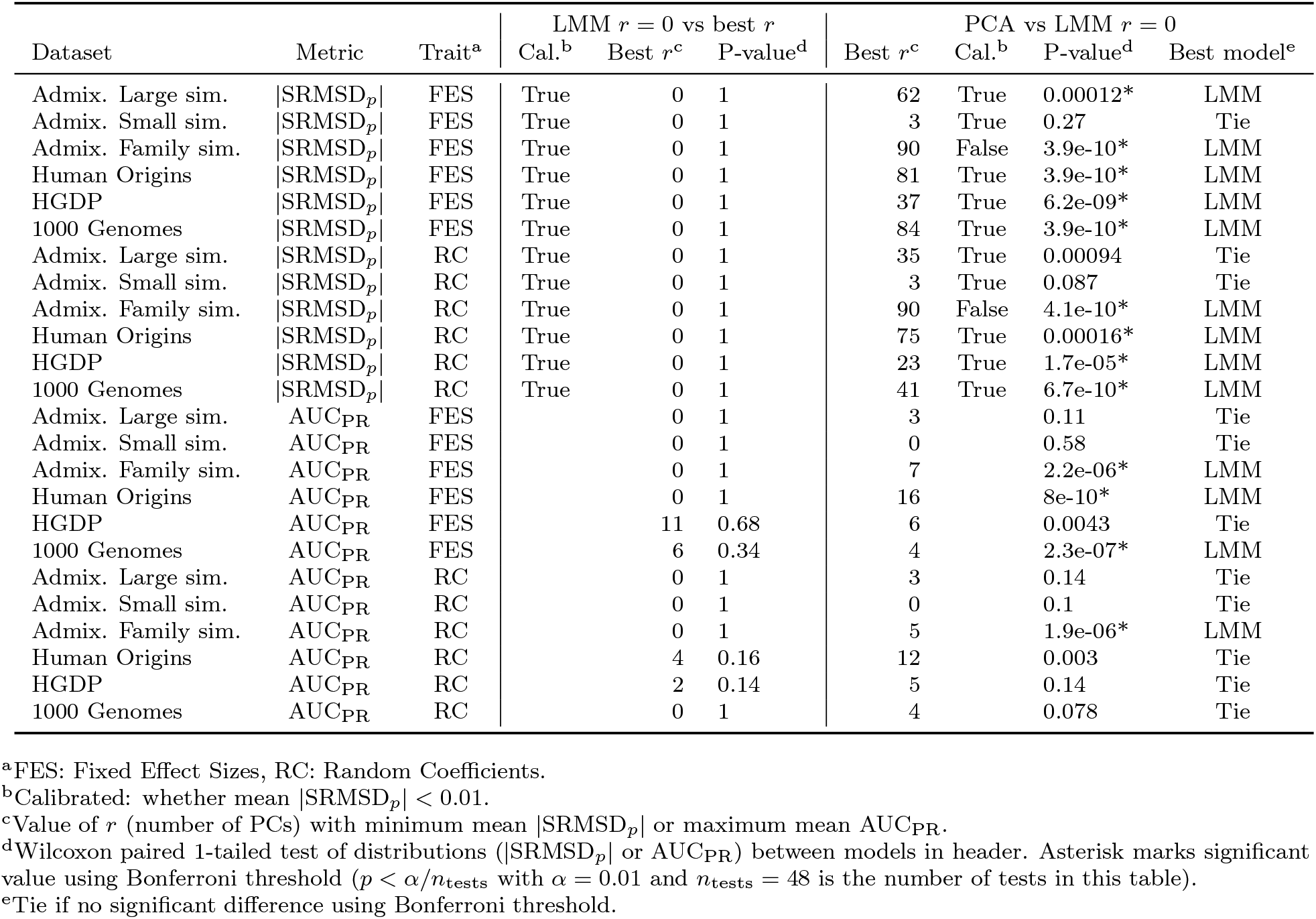
Overview of PCA and LMM evaluations for low heritability simulations.

**Table S3:**
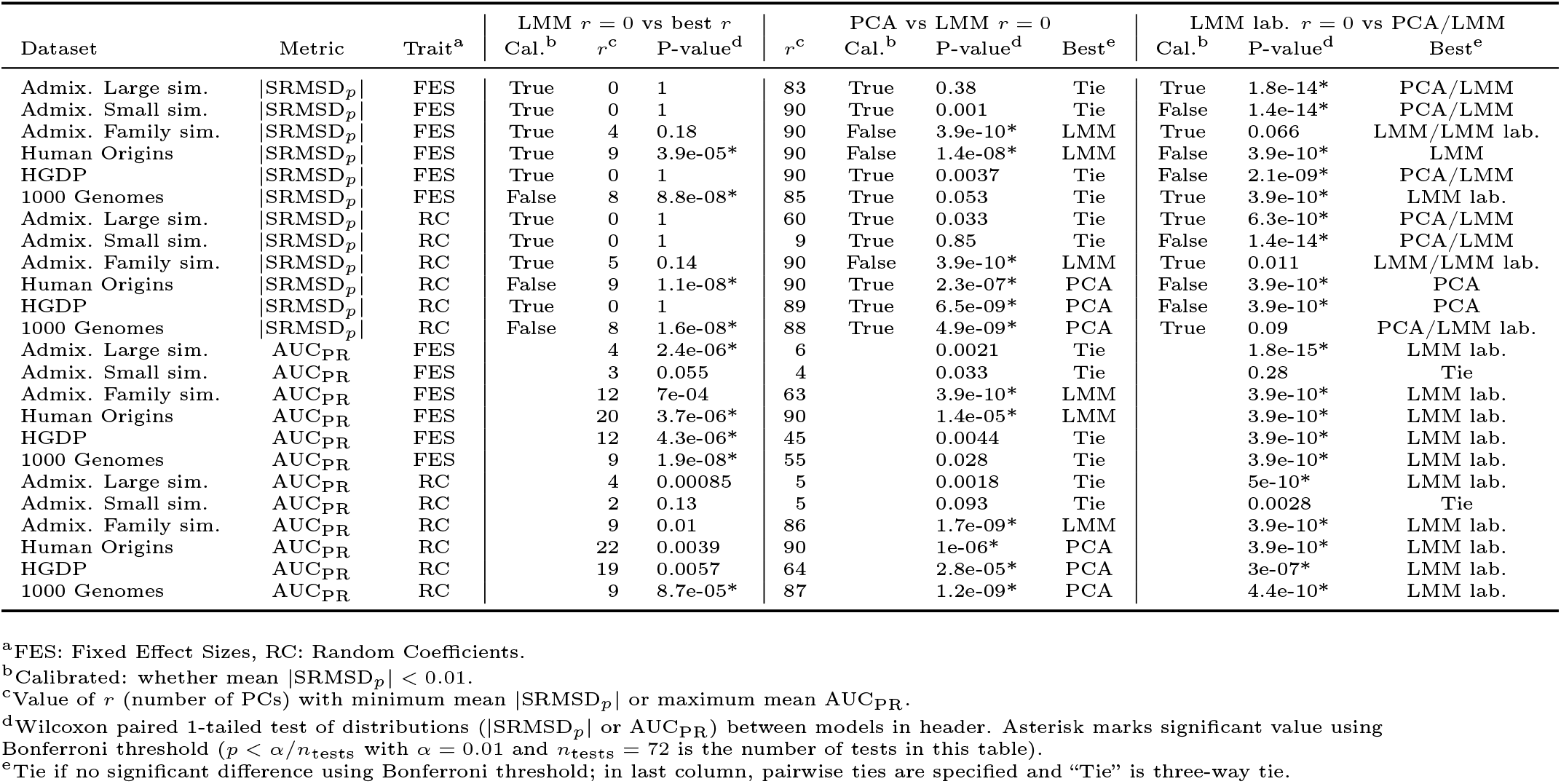
Overview of PCA and LMM evaluations for environment simulations.

## Notes

### Competing Interest Statement

The authors have declared no competing interest.

### Summary of Updates

- We repeated the main evaluations (admixture simulations and real genotypes, both FES and RC trait models) using a lower heritability of h2=0.3, and separately with environment effects (proportion of variance 0.5 for the environment effects, h2=0.3). The low heritability conclusions were the same as for high heritability. The environment evaluations occasionally gave a weak and inconsistent advantage to LMM with PCs or PCA, but directly modeling environment (in an oracle model with the true group labels) yields much greater AUCs compared to indirect modeling via PCs. - We redid the h2=0.8 simulations, which had a small bug discovered after review that caused the residual effects to be incorrectly smaller, so the effective heritability was higher than desired. AUCs are now lower than before, but otherwise all of our conclusions are unchanged. - We now remove loci with more than 10% missingness in HGDP (17% of the loci originally used). For the other real datasets the proportion of such loci was less than 1%, so they were left unfiltered. After redoing all evaluations, our conclusions are unchanged. - Now denote matrix transposition with a prime instead of a T-like symbol that could be confused for the ancestral population T. - We discuss further limitations of our work at greater length, including small sample sizes, exclusion of rare variants, and variations of PCA and LMM. - Numerous minor corrections and clarifications.

